# The BRCA1-A complex restricts replication fork reversal-dependent DNA repair in ATM deficient cells

**DOI:** 10.64898/2026.03.20.713277

**Authors:** Arindam Datta, Jessica Jackson, Yaroslav I Morozov, Jinghan Qiu, Alessandro Vindigni, Roger A. Greenberg

## Abstract

Ataxia Telangiectasia Mutated (ATM) kinase deficiency results in cancer susceptibility and drug sensitivity. Deficiency in either the BRCA1 interacting A complex or XRCC4/Ligase 4 confers resistance to Topoisomerase I or PARP1 inhibitors in ATM-deficient cells. This suggests that BRCA1-A directs toxicity to fork-damaging agents in ATM mutated cells vis-à-vis illegitimate end-joining. Here, we show that ATM inhibition triggers combined SUMO and ubiquitin mediated BRCA1-A damaged fork recognition to restrict end-resection and cause Topoisomerase I inhibitor hypersensitivity. BRCA1-A deficient cells display elevated chromatin accessibility and nuclease activity at damaged forks, coupled with restored resection and drug resistance. Electron microscopy evidence demonstrates that ATM inhibition prevents replication fork reversal, which is restored by BRCA1-A loss to generate substrates for end resection. These findings reveal that BRCA1-A enforces a restrictive chromatin state to suppress the genesis of resection substrates, implicating fork reversal as a key determinant of chemotherapy response in ATM deficient cells.

## Main

Pathogenic mutations to the *BRCA1* or *ATM* genes compromises homologous recombination (HR) and predisposes to breast and ovarian cancers^1–9^. BRCA1 is directly implicated in the multi-step process of HR by promoting end-resection and assembly of the RAD51 filament. Accordingly, BRCA1 mutant cancers display genomic alterations that are characteristic of HR deficiency and either germline or somatic mutations in BRCA1 predict positive responses to PARP inhibitor (PARPi) or platinum-based chemotherapy^9–11^. Moreover, restoration of HR by reversion mutations or loss of 53BP1-Shieldin yields PARPi resistance in BRCA1 mutant cancer cells^11, 12^. Germline ATM deficiency produces profound radiosensitivity, developmental abnormalities, and cancer susceptibility^5^. ATM phosphorylates more than 1000 substrates in response to ionizing radiation (IR), including factors that promote either HR or nonhomologous end-joining (NHEJ)^13^. BRCA1 and many of its interacting partners are confirmed substrates of ATM kinase activity following DNA damage. Despite these commonalities, ATM deficiency differs from loss of BRCA1 in several respects. This includes a different spectrum of cancer susceptibility and developmental abnormalities^5, 14^. These observations are consistent with both shared and distinct aspects of ATM and BRCA1 in response to DNA lesions.

BRCA1 associates stoichiometrically with BARD1 through an amino-terminal interaction^15^. All other BRCA1 interactions are thought to be substoichiometic^16–20^. BRCA1 also interacts with tumor suppressors PALB2 and BRCA2-Rad51 through its coiled coil domain to promote HR^21, 22^. The BRCA1 BRCT repeats are the most common site of cancer associated missense mutations in the BRCA1 gene. The BRCT repeats bind phospho-Serine motifs on Abraxas, Brip1, and CtIP to form the mutually exclusive BRCA1-A, B, and C complexes^17, 23, 24^. Each complex is proposed to direct a different aspect of the damage response. The importance of each interaction with the BRCA1 BRCT repeats remains unknown, however, and interaction defective alleles in either Brip1 or CtIP do not recapitulate loss of function mutation in either the BRCA1 BRCT repeats or Brip1 or CtIP knockouts^25–27^.

The BRCA1-A complex is comprised of BRCA1 in association with ABRAXAS, RAP80, BRCC45, MERIT40, and BRCC36. RAP80 contains tandem ubiquitin interaction motifs (UIMs) at its amino terminus that specifically bind lysine63-linked ubiquitin chains (K63-Ub)^18–20^. The UIM domains are preceded by a SUMO-Interaction motif (SIM) domain. Each motif exhibits a dissociation constant of approximately 20 μM for K63-Ub or SUMO-2, respectively, whereas RAP80 binds to hybrid SUMO2-K63-Ub with 80 fold higher affinity (K_d_<0.2μM)^28^, raising the possibility that hybrid SUMO-ubiquitin chains provide a key damage recognition element for BRCA1-A. BARD1 targets BRCA1 to damage sites by binding nucleosomes that contain unmethylated H4K20 (H4K20me0) and histone H2AK15-Ub^29, 30^. The BRCA1 RING domain interacts with nucleosomes in a ubiquitin-independent manner that allows the BRCA1-BARD1 heterodimer to span di-nucleosomes^31–33^. BARD1 Ankyrin and BRCT interacts with H2AK15Ub, H4K20me0 on one nucleosome and the BRCA1 RING domain interacts with the second nucleosome irrespective of methylation and ubiquitin. How BRCA1-A utilizes these multiple recognition elements provided by RAP80 and BRCA1-BARD1 in the DNA damage response is unknown, as is whether they are individually or collectively required to respond to specific lesions.

Despite extensive investigation into its structural and biochemical properties, little is understood regarding how BRCA1-A contributes to the DNA damage response. BRCA1 damage foci are reduced in A-complex knockout cells but this does not result in similar damage response deficits as observed in context of BRCA1 BRCT mutation^34^. A-complex deficiency is well tolerated in mice, which showed minimal to no changes in comparison to wildtype littermates in response to IR and only modest hypersensitivity to replication fork damaging agents^35, 36^. Moreover, A-complex deficiency increases end resection and HR, indicating that its involvement in BRCA1 damage localization does not promote HR^37, 38^. Recent work speaks to the anti-resection properties of the BRCA1-A complex being a key determinant of chemosensitivity during ATM deficiency. Genome-wide CRISPR-Cas9 screens identified loss of either the A complex or NHEJ as a cause of topoisomerase I inhibitors (TOP1i) and Poly(ADP)ribose polymerase inhibitor (PARPi) resistance specifically in ATM-deficient cancer cells^39^. ATM promotes DNA end resection and HR at replication-associated breaks triggered by TOP1i or PARPi^39–42^. The authors proposed that drug sensitivity in ATM-null cells is due to delayed end resection and HR at broken replication forks, leading to toxic NHEJ ^39, 43^. These results are consistent with prior demonstrations of an anti-resection function of BRCA1-A^37, 38^. A similar synthetic viability interaction occurs upon 53BP1 deletion in BRCA1 mutated cells^44, 45^. However, 53BP1 loss did not confer drug resistance in ATM null cells, suggesting BRCA1-A operates in a distinct manner to prevent resection at damaged forks^39^. This further underscores differences between cancers harboring BRCA1 vs. ATM mutations in the response to chemotherapy, thus representing an important knowledge gap in the treatment of a wide spectrum of cancers.

In this study, we report that BRCA1-A blocks end resection by preventing replication fork reversal in ATM-deficient cells. We show that hypersensitivity to fork damaging agents in ATM inhibited cells requires the complete assembly of the BRCA1-A complex and its combined SUMO and ubiquitin interaction. BRCA1 association with ABRAXAS restricted end-resection and conferred hypersensitivity to Top1i in ATM inhibited cells. Electron microscopy (EM) revealed that ATM kinase activity is required for reversal of damaged replication forks in a manner dependent on fork remodeling proteins, whereas A-complex loss restored fork reversal in ATM inhibited cells commensurate with increased chromatin accessibility, end resection, damage recovery, and Top1i resistance. These findings identify BRCA1-A mediated blockade of replication fork reversal as a critical determinant of end resection and response to cancer therapeutics in ATM deficient cells.

## Results

### BRCA1-A suppresses damaged fork resection in ATM-deficient cells

BRCA1-A loss was reported to confer resistance to fork damaging drugs in ATM-deficient cancer cells^39^. Consistent with the previous results, we observed hypersensitivity of HEK293T cells to nanomolar doses of Topoisomerase 1 inhibitor (Top1i) camptothecin (CPT) when combined with AZD0156^39^, a potent and selective pharmacological ATM kinase inhibitor (ATMi)^42, 43^ (**Fig. 1a** and **Extended Data Fig. 1a, 1b**). ABRAXAS knockout (KO) substantially reduced hypersensitivity to combined CPT and ATMi but did not affect responses to CPT alone. Similar results were obtained in ATM and ABRAXAS double KO HT-29 cells (**Fig. 1b**, and **Extended Data Fig. 1c, 1d**). ABRAXAS knockout also conferred PARPi resistance specifically in the context of ATMi (**Extended Data Fig. 1e**). Restored end resection confers drug resistance in HR-compromised cancer cells^44, 45^. Top1 inhibitors such as CPT generate distinct replication fork intermediates including one-ended double-strand breaks (DSBs) and reversed replication forks. While a high micromolar (µM) dose of CPT induces both one ended DSBs and reversed forks, low nanomolar CPT treatment causes fork reversal without generating DSBs^46, 47^. Reversed fork structures resemble one-ended DSBs and are susceptible to nucleolytic processing by MRE11, EXO1 or DNA2 nucleases^48–51^. Therefore, genetic suppression of hypersensitivity to low nanomolar CPT treatment by BRCA1-A loss in ATM-deficient cells (**Fig. 1a, 1b**) raises the possibility of reversed forks as resection substrates in BRCA1-A deficient cells. We performed RPA immunofluorescence as a read out of single strand DNA generated from end resection in control or ABRAXAS KO HT-29 cells exposed to a low dose (50 nM) of CPT in combination with ATMi (**Fig. 1c, 1d)**. Compared to untreated or only ATMi treated conditions, CPT treatment significantly induced RPA signal in both control and ABRAXAS KO cells. ATMi markedly reduced RPA signal, which was restored in ABRAXAS KO cells, suggesting BRCA1-A restricts end resection upon ATM inhibition. The increased RPA signal was evident in cells that stained positive for EdU incorporation, indicating that elevated resection in A-complex null cells occurs in S-phase (**Fig 1c, 1d**). Similar results were obtained with a high dose (1 µM) of CPT that induces both DSBs and fork reversal (**Extended Data Fig. 2a, 2b)**. We further performed native IdU assay under non-denaturing conditions to detect nascent ssDNA generated specifically at reversed fork structures^52–54^ (**Fig. 1e** and **Extended Data Fig. 2c**). ATMi suppressed nascent ssDNA in cells exposed to a low dose of CPT, while ABRAXAS KO significantly restored it under these conditions. We observed similar results in ATM KO cells with a significant restoration of ssDNA in ATM and ABRAXAS double KO cells (**Fig. 1e** and **Extended Data Fig. 2c**). ATMi did not further exacerbate ssDNA suppression in ATM KO cells, thus attesting to the specificity of the ATM inhibitor. In addition to a high CPT dose (2 µM), BRCA1-A complex loss significantly restored ssDNA in response to as low as 8 nM dose of CPT that does not generally cause DSBs^46, 47^(**Extended Data Fig. 2d)**.

**Fig. 1:**
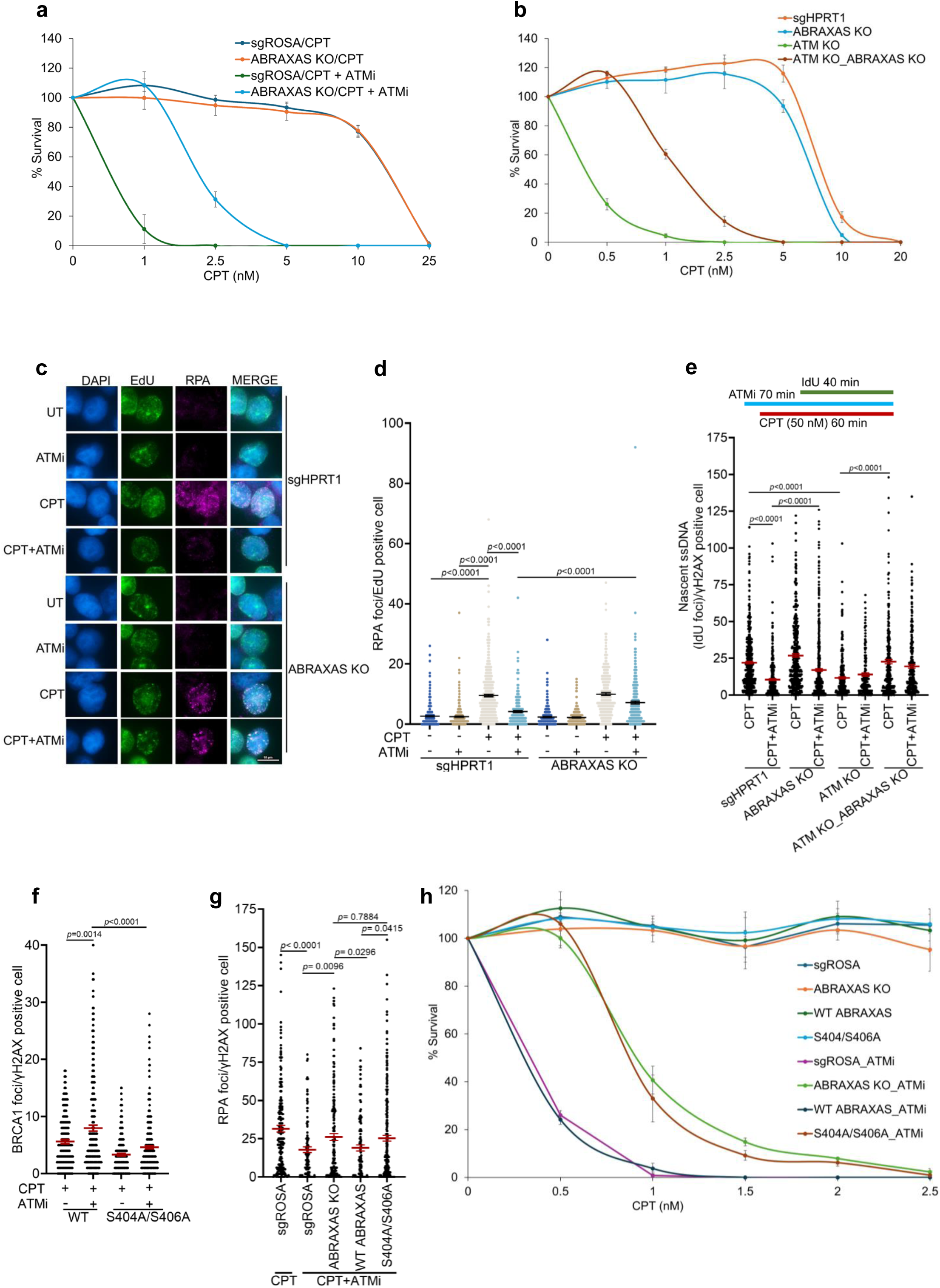
BRCA1 interaction with the A-complex suppresses resection and viability in ATM-deficient cells. **a,** Colony survival of control (sgROSA) and ABRAXAS KO HEK293T cells treated with the indicated doses of CPT alone or in combination with 200 nM ATMi. Relative colony survival (%) in individual genotypes and treatment conditions are shown. Data represent mean ± SD of *n*=3 independent experiments. **b,** Line graph showing relative colony survival of control (sgHPRT1), ABRAXAS KO, ATM KO and ATM-ABRAXAS double KO (DKO) HT-29 cells exposed to the indicated doses of CPT. Data represent mean ± SD of *n*=3 independent experiments. **c,** Representative images of RPA immunofluorescence in control (sgHPRT1) and ABRAXAS KO cells treated with 50 nM CPT +/- 250 nM ATMi for 1 h. Cells were labeled with EdU (10 µM) to identify S-phase cells. Scale bar is shown. **d,** Scatter plot showing significantly reduced RPA foci in CPT-treated EdU positive S-phase cells upon ATMi. ABRAXAS KO significantly restored RPA foci under these conditions. Data represent mean ± SEM derived from n ≥ 200 EdU positive nuclei examined over two independent experiments; *p* values are indicated, unpaired two-tailed t test. **e,** Schematic showing experimental scheme of native IdU assay. Cells were treated with 50 nM CPT +/- 250 nM ATMi as indicated. IdU was added 20 minutes after CPT addition to label newly synthesized DNA encountering CPT-induced lesions. Scatter plot showing quantification of native IdU experiment. Nascent ssDNA (IdU foci) was significantly reduced in either ATM inhibited or ATM KO cells. ABRAXAS KO significantly restored ssDNA under these conditions. Data represent mean ± SEM derived from n ≥ 180 cells examined over two independent experiments; *p* values are indicated, two-tailed Mann–Whitney test. **f,** Scatter plot showing reduced BRCA1 foci in S404A/S406A ABRAXAS expressing cells as compared to WT. WT and S404A/S406A ABRAXAS expressing HT-29 cells were treated with CPT (1 µM) alone or in combination with ATMi (250 nM) for 1 h. ATMi was added 10 min before CPT treatment. Experiments were performed four times with similar results. Data represent mean ± SEM derived from n ≥ 150 cells examined over two independent experiments; *p* values are indicated, unpaired two-tailed t test. **g,** ABRAXAS S404A/S406A mutations restore end resection in ATM-inhibited cells. Scatter plot represents quantification of RPA foci in S404A/S406A cells. Experiments were performed as described in (f) and repeated three times with similar results. Data represent mean ± SEM derived from n ≥ 110 cells examined over two repeats; *p* values are indicated, unpaired two-tailed t test. **h,** Clonogenic survival assay showing CPT+ATMi resistance in HT-29 cells expressing S404A/S406A ABRAXAS. Data are mean ± SD from *n*=3 independent experiments.

### BRCA1 interaction with the A-complex suppresses fork resection and viability in ATM-deficient cells

Suppression of end resection by BRCA1-A complex in ATM-deficient cells led us to further investigate how ATMi triggers BRCA1-A recruitment at damaged forks. The BRCA1-A complex harbors DNA damage recognition moieties contained within RAP80 and BRCA1-BARD1. We therefore examined if both elements were required for its damage localization in ATMi treated cells. Compared to CPT treatment alone, we observed significantly increased RAP80 foci following ATMi (**Extended Data Fig. 3a, 3b)** which is consistent with the previous reports^43^. Similar results were obtained for ABRAXAS and BRCA1 recruitment (**Extended Data Fig. 3c, 3d, 3e** and **3f**), suggesting ATMi triggers BRCA1-A engagement at damaged forks. However, BRCA1 damage localization was significantly impaired in ABRAXAS KO cells (**Extended Data Fig. 3e, 3f**). N-terminal S404 and S406 residues of ABRAXAS (hereafter S404A/S406A) interact with BRCA1 BRCT in a phosphorylation dependent manner^55^. Compared to wild type cells, ATMi-induced BRCA1 damage localization was significantly impaired in cells complemented with S404A/S406A mutant **(Fig. 1f**, and **Extended Data Fig. 4a, 4b**). We also observed a concomitant decrease in ABRAXAS foci formation in these mutant cells (**Extended Data Fig. 4c, 4d**). This indicates that stable chromatin association of the entire BRCA1-A complex with damaged forks requires coordinated A-complex and BRCA1-BARD1 recognition of modified nucleosomes^17, 18, 20, 28–30, 33^.

We further evaluated the effect of complete BRCA1-A assembly on end resection in ATM inhibited cells. While WT ABRAXAS complementation suppressed RPA foci formation in ABRAXAS KO cells, the S404A/S406A mutant did not (**Fig. 1g** and **Extended Data Fig. 4e)**, thereby mimicking A-complex null effects. In agreement, HT-29 cells expressing S404A/S406A mutant showed CPT+ATMi resistance to a similar extent to ABRAXAS KO (**Fig. 1h** and **Extended Data Fig. 5a).** Concordant results were obtained in ABRAXAS KO HEK293T cells complemented with ABRAXAS S404A/S406A **(Extended Data Fig. 5b, 5c).** We conclude that a fully assembled BRCA1-A complex is required to suppress end resection and viability in ATM-deficient cells.

### BRCA1-A regulates chromatin and nuclease accessibility at damaged forks

To identify mechanisms underlying BRCA1-A-mediated suppression of end resection, we characterized the damaged fork proteome in ABRAXAS KO cells using iPOND coupled with mass spectrometry (iPOND-MS). WT or ABRAXAS KO HEK293T cells were exposed to combined CPT and ATMi followed by iPOND purification and mass spectrometry analysis (**Fig. 2a** and **2b**). Western blot revealed specific enrichment of PCNA in iPOND samples as compared to no-click control (**Fig. 2c)**, thereby validating the specificity of the iPOND purification. BRCA1 was depleted in ABRAXAS KO cells, suggesting the A-complex targets BRCA1 to damaged forks (**Fig. 2d)**. MRE11-RAD50-NBS1 (MRN) complex components RAD50 and NBN along with EXO1 nuclease were enriched at damaged forks in ABRAXAS KO cells. Additionally, HR proteins including TIMELESS and TONSL, and several chromatin remodelers were elevated at damaged forks in ABRAXAS KO cells. Gene Ontology (GO) analysis further showed significant enrichment of the DDR response, chromatin organization and remodeling pathways upon BRCA1-A complex loss (**Fig. 2e)**. These results are consistent with a hypothesis that BRCA1-A restricts nuclease accessibility to damaged forks by regulating chromatin state in ATM deficient cells.

**Fig. 2:**
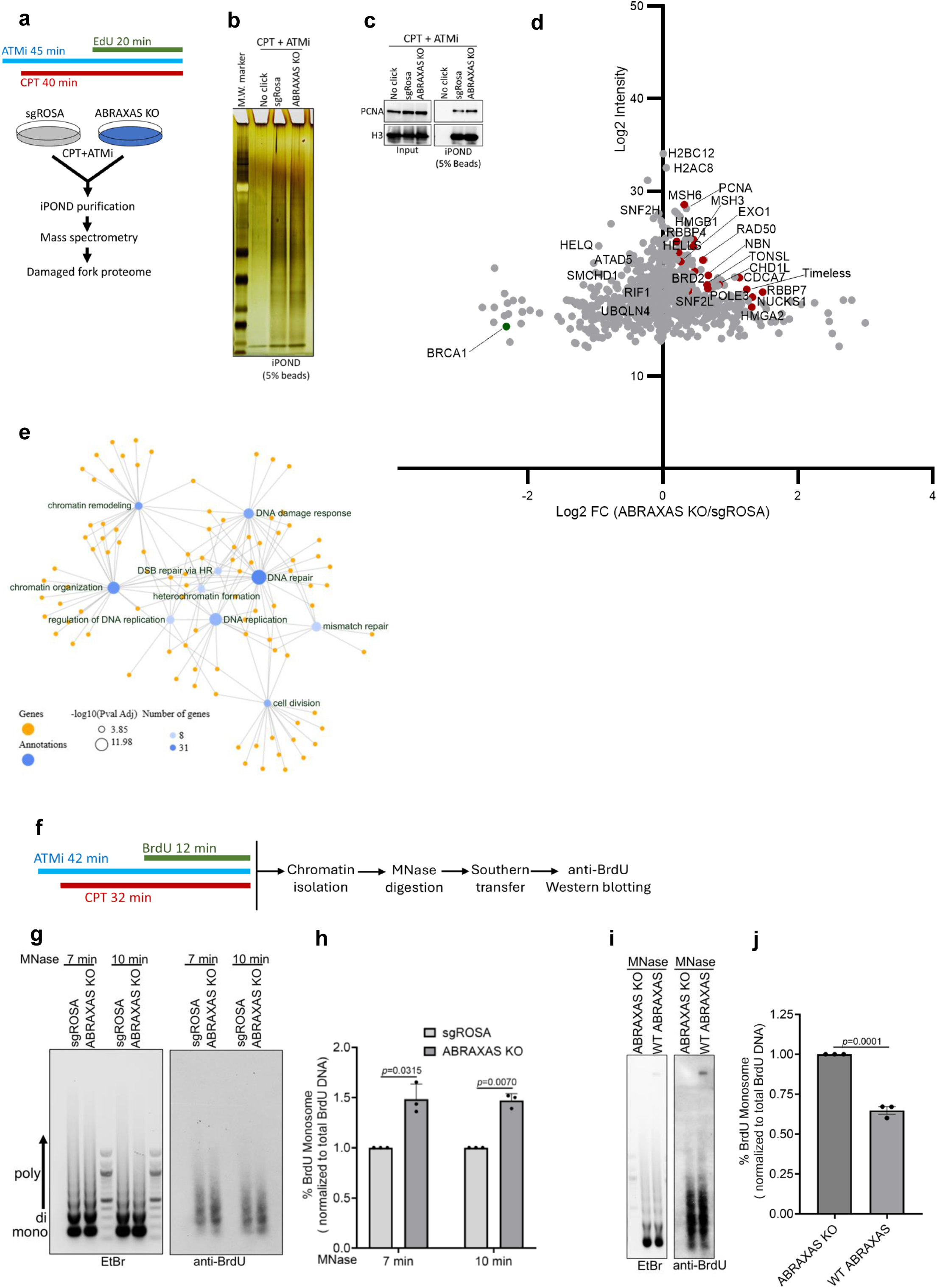
BRCA1-A loss promotes chromatin accessibility and nuclease association at damaged forks in ATM-inhibited cells. **a,** Experimental scheme of the iPOND-MS in control (sgROSA) and ABRAXAS KO cells treated with CPT (1 µM) + ATMi (250 nM). ATMi was added 5 min before CPT treatment and present throughout the experiment. **(b** and **c),** iPOND purified samples (5% beads) were analyzed by silver staining (b) and Western blotting (c). Enrichment of replisome component PCNA in iPOND purified samples but not in control no click indicates specificity of the iPOND purification. **d,** Scatter plot showing iPOND-MS data (*n*=5). X axis shows log2 fold change (FC) of proteins in ABRAXAS KO cells over sgROSA. Y axis shows average log2 intensity of proteins across sgROSA and ABRAXAS KO cells. **e,** Gene-annotation clusters network showing top 10 significantly enriched GO terms for the upregulated proteins in ABRAXAS KO cells. **f,** Experimental scheme of BrdU-coupled chromatin accessibility assay in HEK293T cells treated with CPT (1 µM) +/- ATMi (250 nM). **g,** Representative gel images showing increased MNase accessibility at BrdU-labeled damaged forks in ABRAXAS KO HEK293T cells. Equal number (∼30 X10^6^) of control (sgROSA) and ABRAXAS KO cells were treated with CPT (1 µM) + ATMi (250 nM) and labeled with BrdU. BrdU-labeled nuclei were digested with MNase (250 gel units) for the indicated time followed by chromatin isolation. Equal amount of digested chromatin was run on 1.2% agarose gel (left panel), stained with Ethidium bromide (EtBr) and subsequently transferred to nylon membrane by Southern blotting. MNase accessibility at BrdU labeled damaged chromatin was determined by Western blotting using an anti-BrdU antibody (right panel). **h,** Bar graph showing increased BrdU-labeled mononucleosome fractions in ABRAXAS KO cells as compared to control. Relative percentage of BrdU-labeled mononucleosome fractions normalized to total amount of BrdU-labeled DNA per lane is shown. Data represent mean ± SEM of *n*=3 independent experiments; *p* values are indicated, unpaired two-tailed t test. **(i** and **j),** ABRAXAS complementation in ABRAXAS KO cells reduced MNase accessibility at BrdU-labeled damaged chromatin. Nuclei isolated from ABRAXAS KO and ABRAXAS-complemented HEK2923T cells were treated with CPT (1 µM) + ATMi (250 nM) and were subjected to MNase digestion for 10 min followed by Southern-Western blotting (**i**). Bar graph showing decreased BrdU-labeled mononucleosome fractions upon ABRAXAS complementation (**j**). Data represent mean ± SEM of *n*=3 independent experiments; *p* values are indicated, unpaired two-tailed t test.

Chromatin remodeling allows physical access of nucleases and repair factors to damaged DNA templates^56–61^. ATM kinase activity is critical for chromatin relaxation following DSB induction^62,63^. In response to replication stress, nascent chromatin around replications forks undergoes dynamic changes via nucleosome modifications and remodeling, thereby facilitating fork processing and other transactions^61, 64^. We hypothesized that BRCA1-A regulates chromatin accessibility at damaged forks following ATMi given the increased fork abundance of resection nucleases along with chromatin remodelers in ABRAXAS null cells (**Fig. 2d, 2e**). To determine accessibility specifically at damaged replication forks, we labeled nascent chromatin with a short pulse of BrdU followed by partial MNase digestion and subsequent detection of BrdU labeled chromatin fragments by Southern-Western blotting (**Fig. 2f**). Compared to control cells, we observed significantly increased BrdU-labeled mono nucleosome fractions in ABRAXAS KO cells, suggesting increased chromatin accessibility upon BRCA1-A loss (**Fig. 2g** and **2h**). Complementation of ABRAXAS KO cells with WT ABRAXAS reduced chromatin accessibility at CPT-damaged forks under ATMi conditions (**Fig. 2i** and **2j**), confirming the specificity of these observations.

### Increased end resection upon BRCA1-A loss confers drug resistance in ATM-deficient cells

Controlled nucleolytic processing of damaged forks is critical for HR-dependent fork recovery and cell survival^48, 49, 65^. Because iPOND-MS revealed fork enrichment of EXO1 and MRN nucleases upon BRCA1-A loss, we investigated whether these nucleases promote end resection in this setting. We validated our iPOND-MS results using single molecule PLA-SIRF in cells treated with CPT alone or in combination with ATMi. We observed a significant decrease in the EXO1 PLA-SIRF foci upon ATMi that was restored following ABRAXAS KO, consistent with our iPOND-MS results (**Fig. 3a**, **3b**). We also detected significantly increased fork association of MRE11 nuclease upon BRCA1-A loss in ATMi-treated cells (**Fig. 3c**, **3d**). These results suggest BRCA1-A loss generates replication fork intermediates that engage resection nucleases. We subsequently knocked out MRE11 and analyzed end resection by RPA immunofluorescence in ABRAXAS KO cells treated with CPT plus ATMi (**Fig. 3e, 3f** and **Extended Data Fig. 6a**). MRE11 KO significantly reduced RPA foci in ABRAXAS KO cells exposed to combined CPT and ATMi (**Fig. 3f**). We also observed a partial reduction in RPA foci formation following EXO1 KO in ABRAXAS-deficient cells (**Fig. 3g, 3h** and **Extended Data Fig. 6b**), indicating enhanced end resection upon BRCA1-A loss in ATM-deficient cells is dependent on MRE11 and EXO1 nucleases. Consistent with the end resection data, EXO1 KO partially resensitized ABRAXAS KO cells to combined CPT and ATMi treatment (**Fig. 3i** and **Extended Data Fig. 6c**). Together, these results suggest that resistance to combined Top1i and ATMi treatment upon BRCA1-A loss is attributed to MRE11 and EXO1-mediated end resection.

**Fig. 3:**
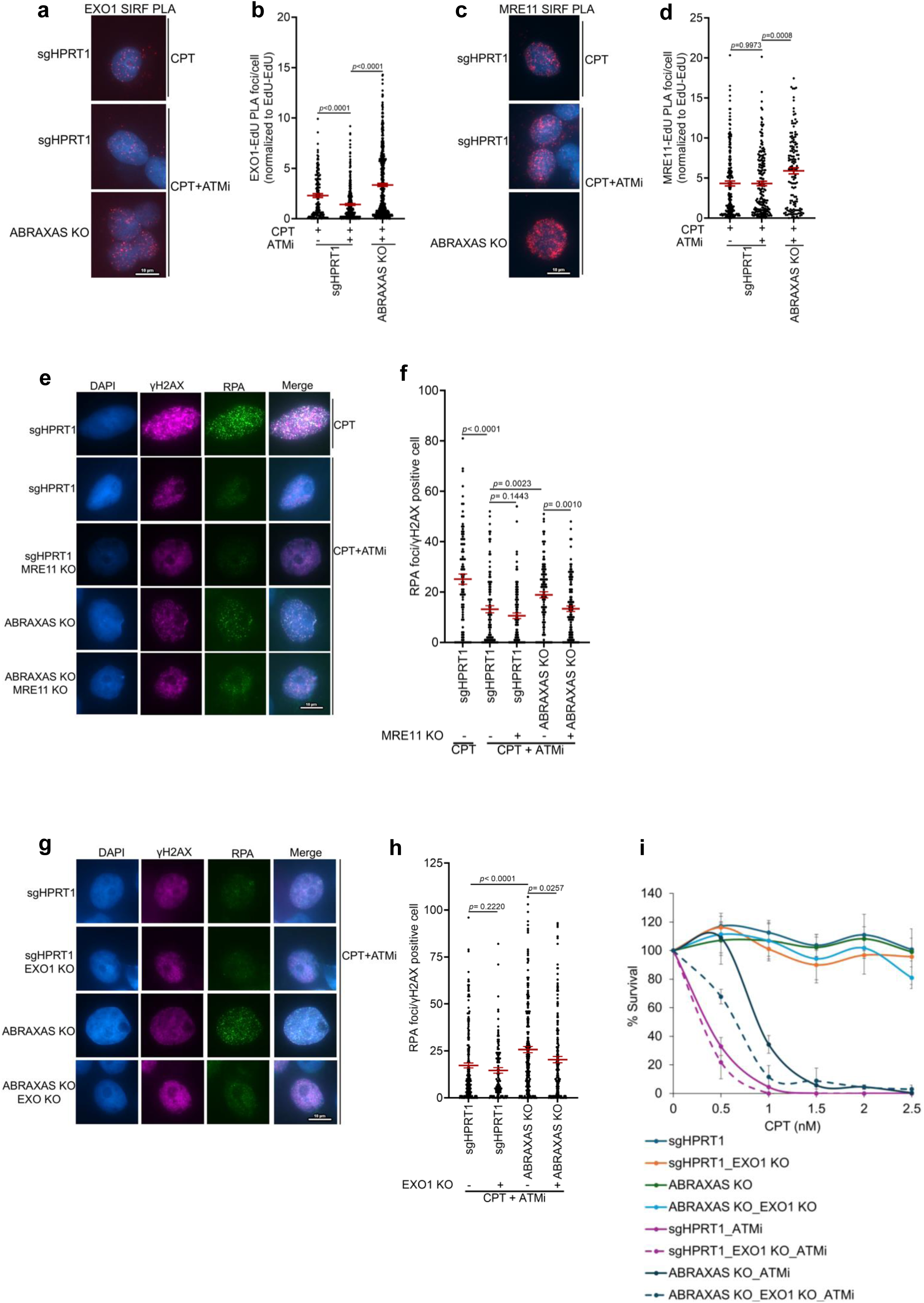
EXO1 and MRE11-mediated end resection confers drug resistance in BRCA1-A deficient cells. **(a and b),** Representative images of PLA-SIRF experiments showing EXO1 binding to replication forks in control (sgHPRT1) and ABRAXAS KO HT-29 cells treated with CPT (1 µM) +/- ATMi (250 nM) for 40 min. Cells were optionally pretreated with ATMi for 10 min and labeled with EdU during the last 20 min of CPT treatment. ATMi was present throughout the experiment. Images were acquired with 100X magnification. Scale bar is shown (**a**). Scatter plot showing quantification of EXO1-EdU SIRF foci normalized to average EdU-EdU signal in each experimental condition (**b**). Data represent mean ± SEM derived from n ≥ 139 cells examined over two independent experiments; *p* values are indicated, unpaired two-tailed t test. **(c** and **d),** PLA-SIRF experiments showing increased MRE11 fork binding in ABRAXAS KO cells treated with CPT and ATMi as described in (a). Representative images captured with 100X magnification are shown. Scale bar is shown (**c**). Scatter plot showing quantification of the MRE11-EdU SIRF foci normalized to average EdU-EdU signal in individual experimental conditions (**d**). Data represent mean ± SEM derived from n ≥ 140 cells examined over two independent experiments; *p* values are indicated, unpaired two-tailed t test. **(e and f**) Representative images of RPA immunofluorescence in the indicated genotypes following treatment with CPT (1 µM) +/- ATMi (250 nM) for 1 h. ATMi was added 10 min before CPT treatment. γH2AX marks cells with CPT-induced replication damage. Scale bar is shown (**e**). Scatter plot showing quantification of the RPA foci (**f**). Data represent mean ± SEM derived from n ≥ 90 cells examined over two independent experiments; *p* values are indicated, unpaired two-tailed t test. **(g** and **h),** Representative images of RPA immunofluorescence in the indicated genotypes treated with combined CPT (1 µM) and ATMi (250 nM) for 1 h. ATMi was added 10 min before CPT treatment. Scale bar is shown (**g**). Scatter plot showing quantification of the RPA immunofluorescence (**h**). Data represent mean ± SEM derived from n ≥ 100 cells examined over two biological repeats; *p* values are indicated, unpaired two-tailed t test. **i,** Colony survival assay showing EXO1 KO resensitizes ABRAXAS KO cells to combined CPT and ATMi treatment. Data represent mean ± SD derived from *n*=3 independent experiments.

### ATM inhibition promotes SUMO and ubiquitin dependent BRCA1-A localization to damaged forks

Double strand break (DSB) recognition by the A-complex is mediated by RAP80 binding to K63-linked polyubiquitin chains via its tandem ubiquitin-interacting motifs (UIMs)^18–20^. Additionally, RAP80 harbors a small SUMO interaction motif (SIM) that contributes to RAP80 and BRCA1 damage localization^28^ (**Fig. 4a**). *In vitro* binding experiments demonstrated that recombinant RAP80 has an 80-fold higher binding affinity for hybrid SUMO-ubiquitin chains compared to monomeric SUMO-2 or diUb alone^28^. We therefore assessed how RAP80 ubiquitin and SUMO recognition contributes to BRCA1-A damage accumulation during ATMi. To this end, we complemented RAP80 KO cells with RAP80 WT or deletion mutants lacking UIMs, SIM, or both SIM and UIM domains (**Fig. 4b**) and tested RAP80 damage localization. WT cells showed increased RAP80 damage accumulation following combined CPT and ATMi treatment (**Fig. 4c** and **Extended Data Fig. 7a**). In contrast, individual deletion of UIM or SIM showed a significant decrease in RAP80 foci formation which was even more pronounced upon combined UIM and SIM loss (**Fig. 4c** and **Extended Data Fig. 7a)**. While deleting the UIMs alone did not affect BRCA1 recruitment, loss of the SIM domain reduced BRCA1 foci and combined SIM and UIM deletion suppressed BRCA1 damage localization further (**Fig. 4d** and **Extended Data Fig. 7b**). In accordance, inhibition of SUMOylation by the SUMO E1 inhibitor ML-792 significantly reduced RAP80-EdU PLA signal (**Extended Data Fig. 7c, 7d**). Consistent with these results, SIRF-PLA experiments revealed a significant increase in SUMO 2/3 abundance at forks upon ATMi (**Extended Data Fig. 7e**). Ubiquitin signaling at damaged forks was also found to be elevated upon ATMi (**Extended Data Fig. 7f**). To determine the impact of ubiquitin or SUMO recognition on fork resection, we examined RPA foci formation in RAP80 WT or mutant cells exposed to CPT plus ATMi. WT RAP80 complementation suppressed end resection in RAP80 KO cells, while loss of the SIM domain alone, but not the UIM, partially restored it (**Fig 4e** and **Extended Data Fig. 8a**). A more significant increase in RPA foci over WT cells occurred upon concurrent deletion of SIM and UIM. This reflects requirements for both SUMO and ubiquitination at damaged forks upon ATMi. This led us to further investigate whether SIM and UIM deletion affects CPT sensitivity in ATM-inhibited cells. RAP80 KO showed CPT resistance when combined with ATMi, whereas complementation with WT RAP80 reinstated drug sensitivity (**Fig. 4f** and **Extended Data Fig. 8b**). While deletion of SIM domain exhibited partial resistance as compared to WT cells, UIM deletion did not. Consistent with the end resection data, we observed a more significant increase in drug resistance upon combined deletion of SIM and UIM domains. Together, these results indicate that SUMO is the primary determinant for BRCA1-A damaged fork response, and that combined SUMO and ubiquitin recognition by RAP80 is required for maximal suppression of resection and drug hypersensitivity in ATM-deficient cells.

**Fig. 4:**
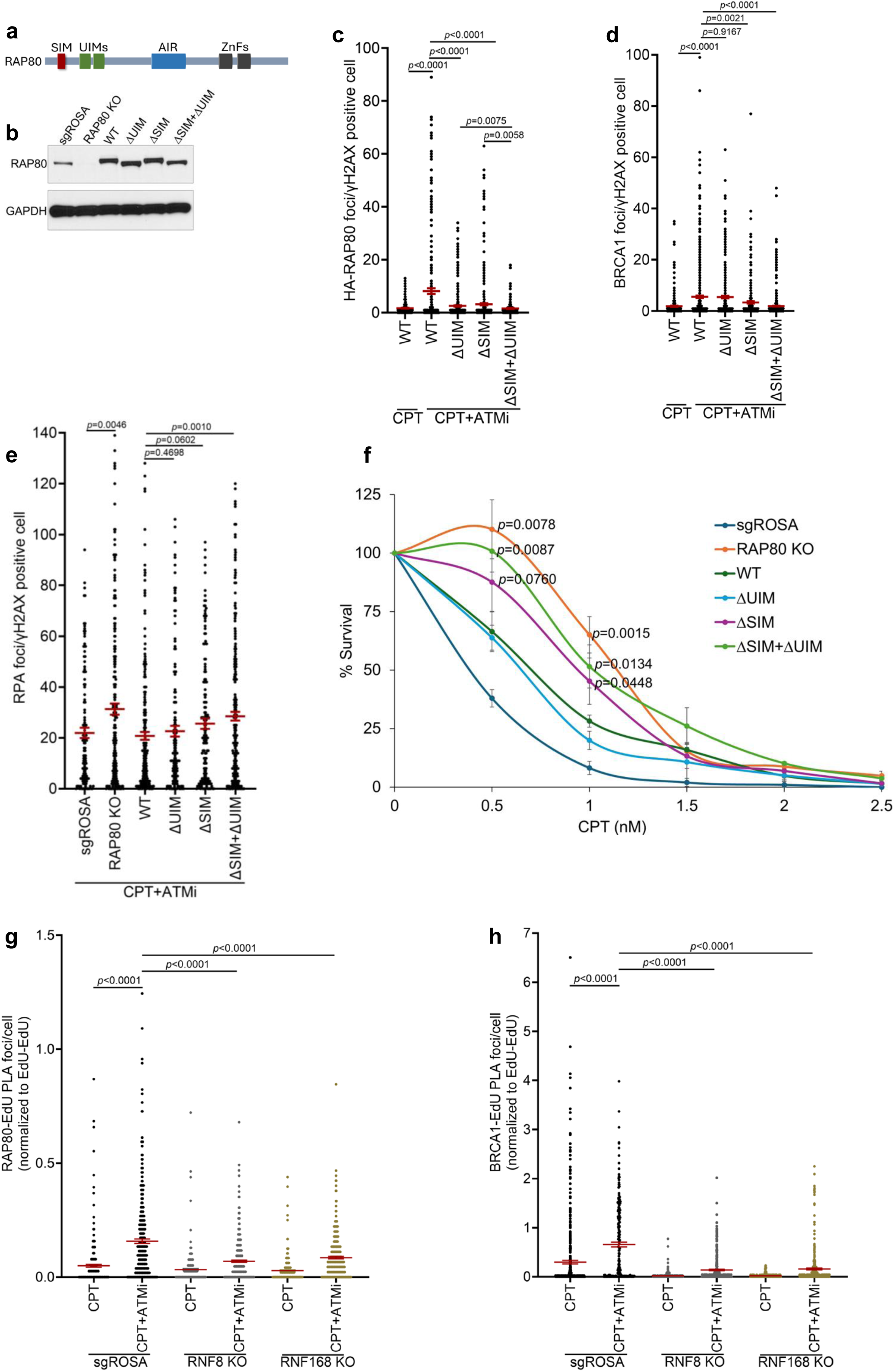
Combined SUMO-ubiquitin recognition determines BRCA1-A function at damaged forks. **a**, Schematic showing structural domains of RAP80. SIM, SUMO interaction motif; UIMs, Ubiquitin interaction motifs; AIR, ABRAXAS interacting region; ZnFs, Zinc finger domains. **b**, Western blots showing RAP80 protein levels in RAP80 KO HT-29 cells complemented with FLAG-HA tagged WT, ΔUIM, ΔSIM or ΔSIM + ΔUIM RAP80. GAPDH serves as loading control. **c,** Scatter plot showing quantification of HA-RAP80 foci in WT, ΔUIM, ΔSIM or ΔSIM + ΔUIM expressing HT-29 cells treated with CPT (1 µM) +/- ATMi (250 nM) for 1 h. ATMi was added 10 min before CPT treatment. Data represent mean ± SEM derived from n ≥ 230 cells examined over two independent experiments; *p* values are indicated, unpaired two-tailed t test. **d,** Scatter plot showing quantification of BRCA1 foci in the genotypes as indicated in (c). Data are mean ± SEM from n ≥ 280 cells examined over two repeats; *p* values are indicated, unpaired two-tailed t test. **e,** Quantification of RPA immunofluorescence in HT-29 cells expressing RAP80 WT or the indicated deletion mutants upon treatment with CPT (1 µM) + ATMi (250 nM) for 1 h. Data represent mean ± SEM derived from n ≥ 125 cells examined over two repeats; *p* values are indicated, unpaired two-tailed test. **f,** Line graph showing relative colony survival of control (sgROSA) and RAP80 KO HT-29 cells harboring empty vector (EV) or complemented with WT or RAP80 deletion mutants. Cells were treated with 100 nM ATMi and the indicated doses of CPT. *p* values were determined for differences in colony survival for RAP80 KO (EV), ΔSIM or ΔSIM + ΔUIM harboring cells over WT at CPT concentrations of 0.5 and 1 nM; unpaired two-tailed t test. Data represents mean ± SD from *n*=3 biological experiments. **g,** Scatter plot showing quantification of RAP80-EdU PLA in control (sgROSA), RNF8 or RNF168 KO HT-29 cells treated with 50 nM CPT +/- ATMi (250 nM). Cells were optionally pretreated with ATMi for 10 minutes followed by CPT treatment for 40 minutes. EdU was added during the last 20 min of CPT treatment. ATMi was present throughout the experiment. Data represent mean ± SEM derived from n ≥ 279 nuclei examined over two independent experiments; *p* values are indicated, unpaired two-tailed t test. **h,** Quantification of BRCA1-EdU PLA as described in (e). Data are mean ± SEM from n ≥ 239 nuclei pooled from two independent experiments; *p* values are indicated, unpaired two-tailed t test.

Given their central role in DSB ubiquitin signaling^66^, we next tested if E3 ubiquitin ligases RNF8 and RNF168 are involved in fork ubiquitylation. Fork ubiquitylation was significantly diminished upon RNF8 or RNF168 KO in ATMi-treated cells (**Extended Data Fig. 9a, 9b**), accompanied by reduced fork binding of RAP80 (**Fig. 4g**) and BRCA1 (**Fig. 4h**). These results were further supported by increased fork binding of RNF8 (**Extended Data Fig. 9c**) and RNF168 (**Extended Data Fig. 9d**) upon ATMi, consistent with previous findings^43^. ATM-dependent phosphorylation of H2AX and MDC1 promotes RNF8 and RNF168 engagement on damaged chromatin^66^. We found that H2AX or MDC1 KO led to reduced RNF8-/RNF168-EdU PLA signal compared to control cells (**Extended Data Fig. 9c, 9d**). However, RNF8 or RNF168 fork abundance was found to be increased following ATMi even in H2AX or MDC1 KO cells (**Extended Data Fig. 9c, 9d**), potentially reflecting an H2AX/MDC1 independent mechanism of RNF8/168 fork recruitment under ATM-deficient condition.

### BRCA1-A prevents fork reversal in ATM-deficient cells

Top1 poisons induce high rates of replication fork reversal, a RAD51-dependent protective mechanism that limits formation of DSBs and facilitates repair or bypass of DNA lesions^46, 47, 67^. Fork reversal is catalyzed by ATP-dependent DNA translocases including SMARCAL1, ZRANB3, HLTF and FBH1^68^. Immunofluorescence imaging results of RPA (**Fig 1c, 1d**) and native IdU (**Fig. 1e**, and **Extended Data Fig. 2c, 2d**) using low nanomolar dose of CPT are consistent with increased end-resection at reversed fork substrates in BRCA1-A deficient cells. We extended this analysis in cells treated with hydroxyurea (HU), which promotes degradation of reversed forks in BRCA1 or BRCA2 deficient cells. BRCA1 depletion led to fork degradation as expected, while ABRAXAS KO did not cause degradation of HU-stalled forks (**Extended Data Fig. 10a**). We observed a moderate but significant decrease in replication tract length in ABRAXAS KO cells when exposed to ATMi (**Extended Data Fig. 10a**). Similar results were observed in native IdU immunofluorescence experiments (**Extended Data Fig. 10b**), suggesting controlled resection of HU-induced reversed forks occurs when BRCA1-A loss is concurrent with ATM-deficiency.

We performed electron microscopy (EM) to quantitatively determine the abundance of reversed fork structures following a 50 nM dose of CPT treatment that is established to cause fork reversal without generating DSBs^46, 47, 69, 70^ (**Fig. 5a, 5b**). Consistent with published results^46, 47^, we detected an average of 32% reversed fork structures upon CPT treatment across three independent biological experiments (**Fig. 5b**). We observed a > 3-fold decrease in fork reversal events in ATMi + CPT treated cells, revealing a critical role of ATM kinase in promoting damaged fork reversal. Notably, ABRAXAS KO restored fork reversal by 2.5-fold in ATMi treated cells (**Fig. 5b**). The frequency of post-replicative ssDNA gaps (**Fig. 5c)** and fork junction gaps (**Fig. 5d**) were increased upon ATMi. This may reflect accumulation of uncoupled forks with extended parental ssDNA at fork junctions and repriming, respectively due to impaired fork reversal in ATM-inhibited cells. ABRAXAS KO did not alter the frequency of gap formation (**Fig. 5c, 5d**), suggesting that this lesion type is not responsible for the associated drug resistance.

**Fig. 5:**
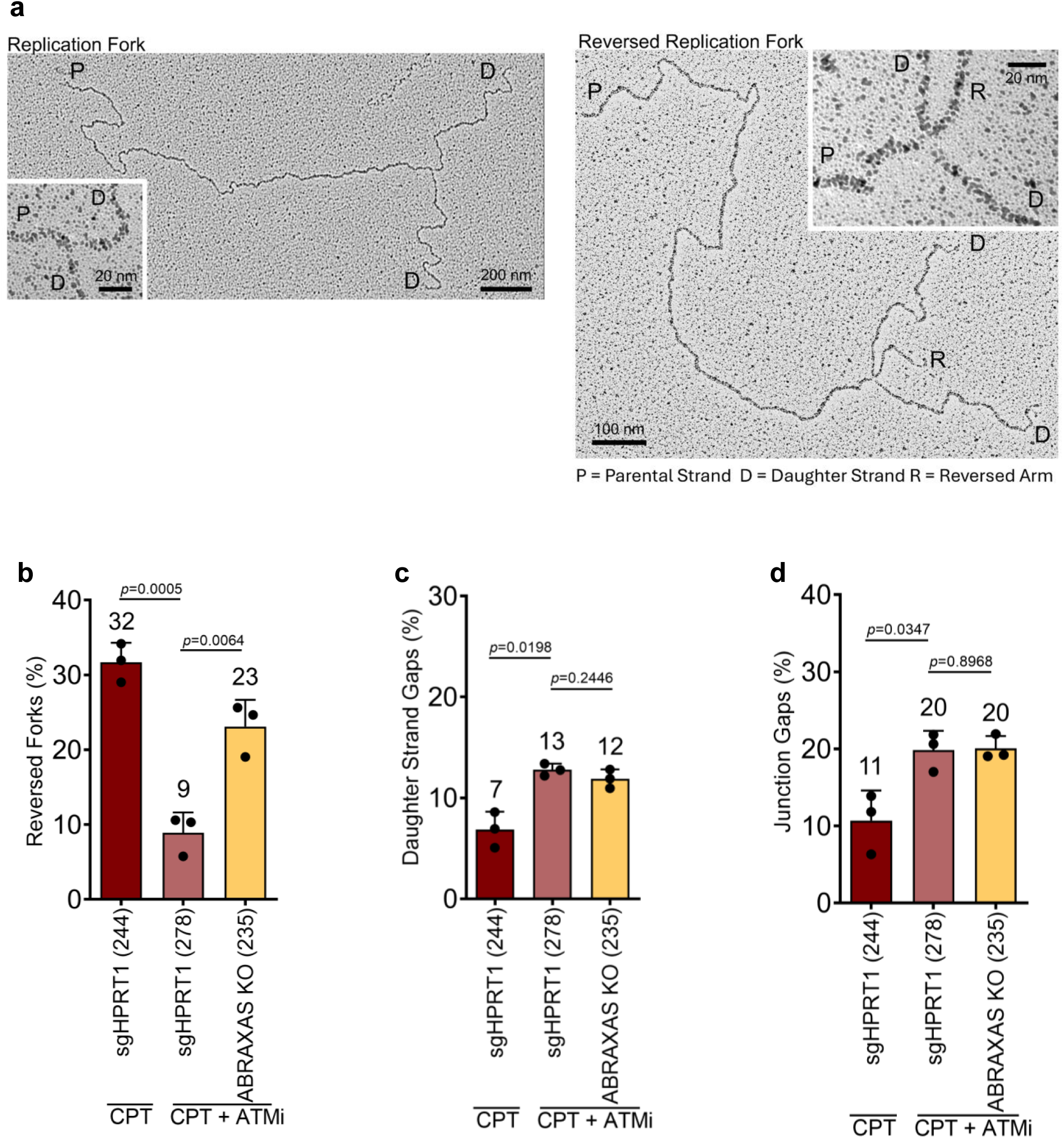
BRCA1-A complex prevents fork reversal upon ATM inhibition. **a,** Left panel: Electron micrograph showing a typical replication fork with a three-way junction. Right panel: Electron micrograph of a typical reversed replication fork with a four-way junction. Representative images are from psoralene-crosslinked genomic DNA isolated from ABRAXAS KO HT-29 cells treated with CPT+ATMi. P, parental strands; D, daughter strands; R, reversed arm. Cells were treated with 50 nM CPT for 1 h. ATMi (250 nM) was added optionally 10 min prior to CPT treatment and present throughout the experiment. **b,** Bar graph showing frequency of fork reversal in control (sgHPRT1) cells treated with CPT+/-ATMi or ABRAXAS KO cells treated with CPT+ATMi. Fractions of reversed forks (number above bars) as a percentage of total number of replication intermediates analyzed (bottom, within parentheses) are shown. **(c** and **d),** Bar graphs showing fractions of post-replicative daughter strand gaps (**c**) and junction gaps (**d**) detected under experimental conditions as described in (a). In (b), (c) and (d), data represent mean ± SD from *n*=3 independent biological experiments; *p* values are indicated; unpaired two-tailed t test.

### Reversed forks are resection substrates in cells deficient for ATM and BRCA1-A

Because CPT-induced fork reversal slows down replication, we assessed fork speed using DNA fiber analysis as a surrogate measure of fork reversal ^47, 67, 69, 71^. ATMi considerably rescued replication tract length in CPT-treated cells, consistent with compromised fork reversal under this condition (**Fig. 6a**). However, ATMi did not significantly increase fork speed in ABRAXAS KO cells, thus reflecting restored fork reversal in these cells (**Fig. 6a**), in agreement with EM data (**Fig. 5**).

**Fig. 6.**
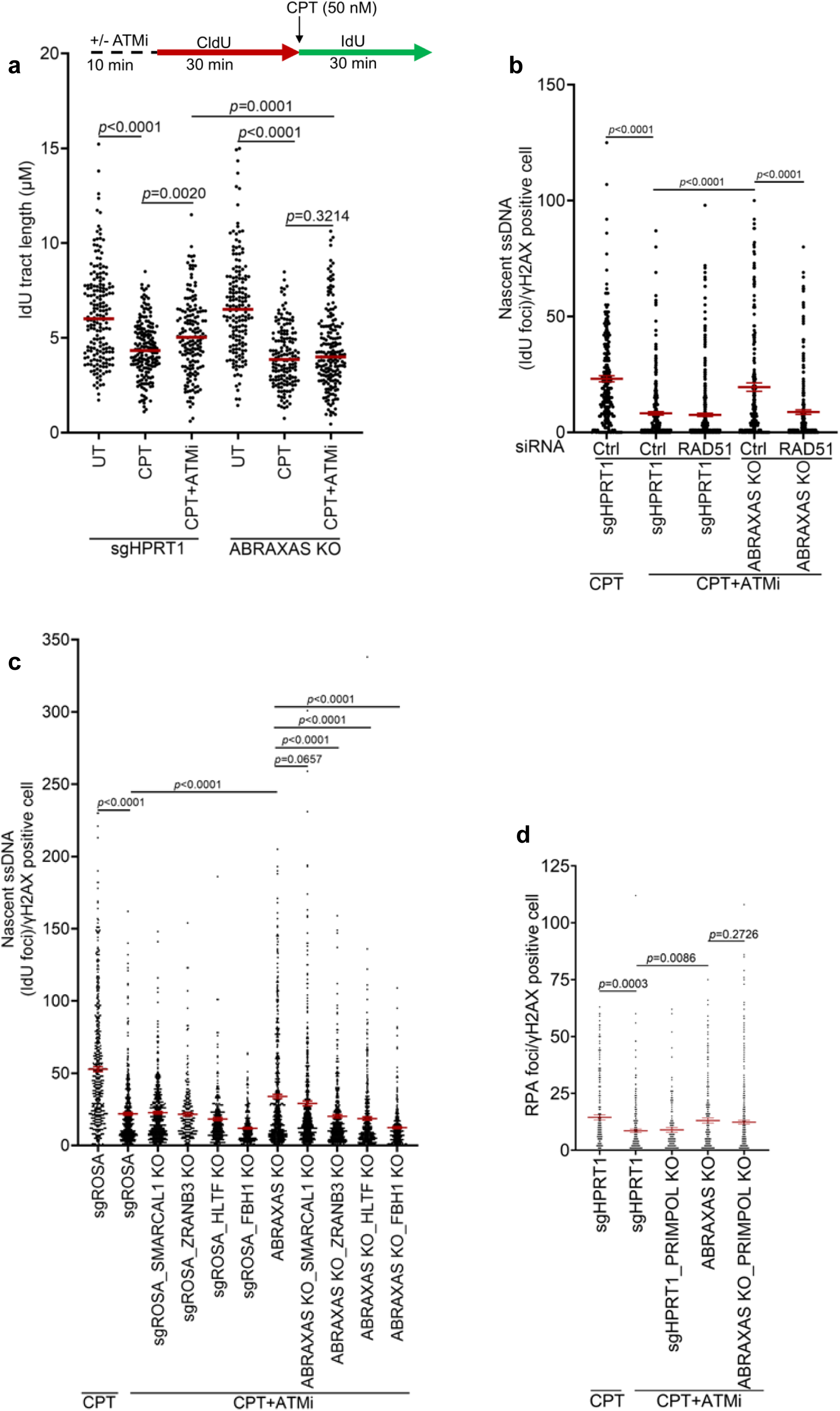
BRCA1-A blocks resection at reversed forks in ATM-inhibited cells. **a,** Schematic showing the strategy of the DNA fiber assay to examine CPT-induced fork slowing as a read out of fork reversal. Control (sgHPRT1) or ABRAXAS KO HT-29 cells were optionally pre-treated with ATMi (250 nM) for 10 min before CldU labeling. ATMi was present throughout the experiment and cells were treated with 50 nM CPT during the IdU labeling. Scatter plot represents IdU tract length in individual genotypes treated as indicated. Data were derived from n ≥ 150 DNA fibers pooled from two independent experiments. Horizontal red bars indicate median IdU tract length; two-tailed Mann–Whitney test. **b,** Scatter plot showing significantly reduced nascent ssDNA (IdU foci) in ABRAXAS KO cells upon RAD51 depletion. Data represent mean ± SEM derived from n ≥ 230 cells examined over two independent experiments; *p* values are indicated, two-tailed Mann–Whitney test. **c,** Quantification of ssDNA (IdU foci) in U2OS control (sgROSA) and ABRAXAS KO cells upon SMARCAL1, ZRANB3, HLTF or FBH1 KO. Cells were treated with 50 nM CPT for 60 minutes. IdU was added during the last 40 minutes of CPT treatment. Cells were optionally pre-treated with ATMi (250 nM) 10 minutes before CPT addition and continued throughout the experiment. Data represent mean ± SEM derived from n ≥ 272 cells examined over two independent experiments; *p* values are indicated, two-tailed Mann–Whitney test. **d,** Quantification of RPA immunofluorescence in control (sgHPRT1) and ABRAXAS KO HT-29 cells upon PRIMPOL KO. Cells were treated with 50 nM CPT +/- ATMi (250 nM) as indicated for 1 h. Experiments were performed three times with similar results. Data represent mean ± SEM derived from n ≥ 129 cells examined over two independent experiments; *p* values are indicated, two-tailed Mann–Whitney test.

These results imply that BRCA1-A loss generates substrates for nucleolytic resection by restoring fork reversal in ATM-inhibited cells. We treated ABRAXAS KO cells with 50 nM CPT +/- ATMi and examined nascent ssDNA by native IdU assay after depletion of RAD51, which is reported to be essential for replication fork reversal^47, 67^. RAD51 depletion diminished nascent ssDNA in ABRAXAS KO cells exposed to ATMi (**Fig 6b** and **Extended Data Fig. 11a, 11b**), consistent with fork reversal being the source of resection substrates in BRCA1-A deficient cells. To further test the importance of fork reversal for resection, we determined ssDNA levels upon knockout of fork reversal enzymes SMARCAL1, ZRANB3, HLTF or FBH1 (**Fig. 6c** and **Extended Data Fig. 11c**). SMARCAL1 KO did not significantly change IdU labeled ssDNA in ABRAXAS-deficient cells. In contrast, KO of HLTF, ZRANB3 or FBH1 suppressed ssDNA in ABRAXAS KO cells exposed to CPT+ATMi, with FBH1 loss showing a dramatic reduction in native ssD

NA generation (**Fig. 6c** and **Extended Data Fig. 11c**). Post-replicative single-stranded DNA gaps also arise following replication stalling via PRIMPOL mediated repriming ^71–73^, representing a second possibility for DNA substrates that could serve as resection substrates in the context of concomitant ATM kinase and BRCA deficiency. Nucleolytic processing would further extend these gaps resulting in RPA coated ssDNA on parental DNA strand^74, 75^. However, PRIMPOL KO did not affect RPA foci in ABRAXAS KO cells treated with 50 nM CPT +/- ATMi, suggesting resection at gaps is not responsible for the elevated ssDNA (**Fig. 6d** and **Extended Data Fig. 11d, 11e**).

### BRCA1-A loss restores fork restart and chromosome stability in ATM-deficient cells

Fork reversal allows damage fork recovery by homology-directed repair (HDR). Controlled nucleolytic resection of a reversed fork generates 3′-end required for Rad51-mediated strand invasion of the duplex ahead of the fork to promote replication restart^49, 76^. We investigated how ATM and BRCA1-A status affect fork restart following CPT treatment using the single molecule DNA fiber technique (**Fig. 7a**). While ATMi significantly decreased fork restart, ABRAXAS KO partially restored it (**Fig. 7b**). We further determined DNA fiber tract (IdU) length following replication restart as a read out of fork restart efficiency (**Fig. 7c** and **Extended Data Fig. 12a, 12b)**. We observed a rescue of restarted tract length upon ABRAXAS KO, suggesting BRCA1-A complex loss allows efficient replication resumption of CPT-stalled forks in ATMi treated cells.

**Fig. 7:**
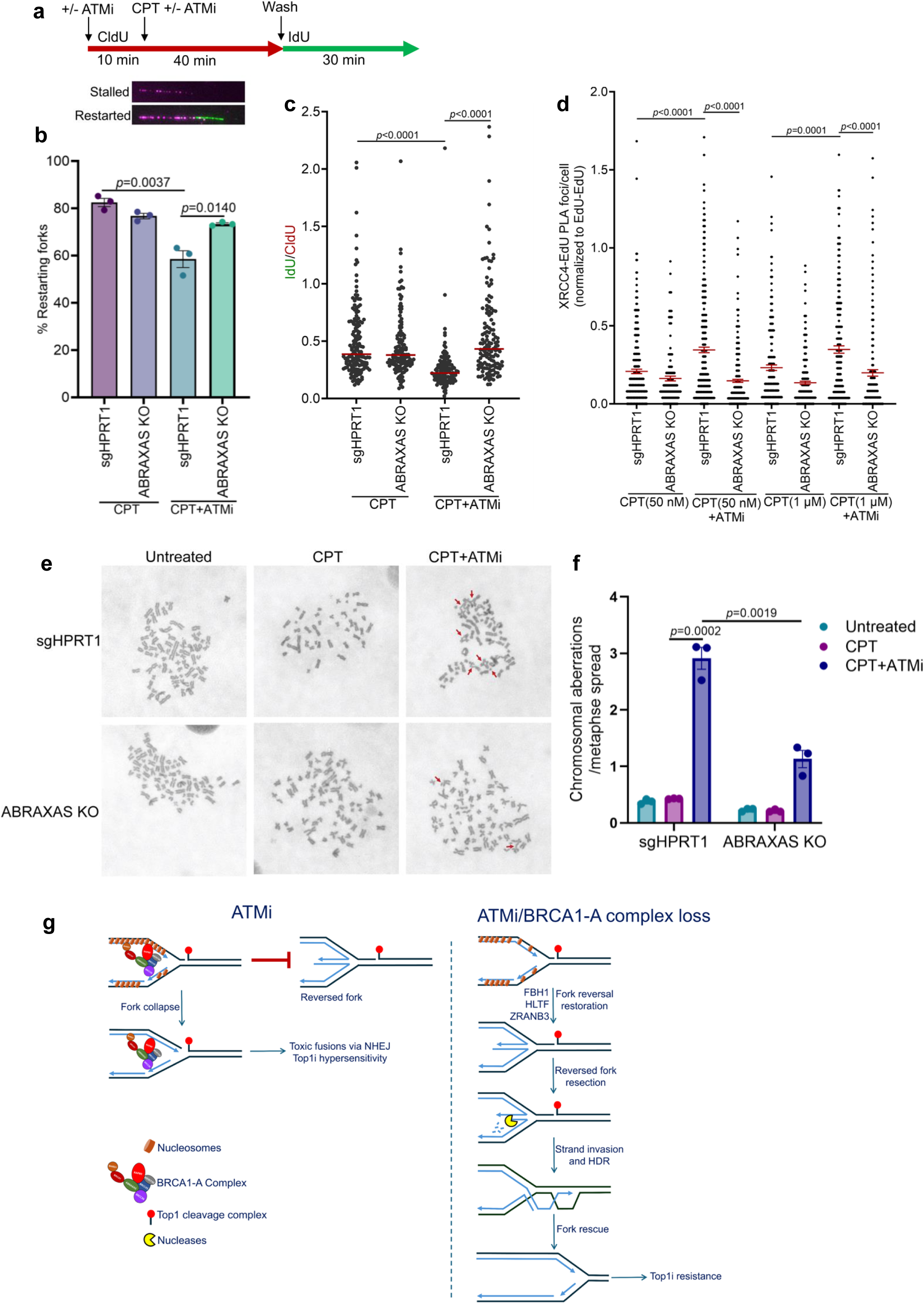
BRCA1-A loss promotes damaged fork recovery and suppresses chromosomal aberrations in ATM-inhibited cells. **a,** Schematic of DNA fiber assay to determine fork restart following CPT (50 nM) +/- ATMi (250 nM) treatment (upper panel). Representative fiber images of a stalled and a restarted fork are shown (lower panel). **b,** Bar graph showing fractions of restarted forks (red-green) as a percentage of total red labeled fibers examined under each experimental condition. More than 300 fibers were analyzed over *n*=3 repeats. Data represent mean ± SEM; *p* values are indicated, unpaired two-tailed t test. **c,** Scatter plot showing IdU/CldU ratios determined from the experiment as described in (a). *p*-values were determined from n ≥ 135 DNA fibers using two-tailed Mann–Whitney test. **d,** Quantification of XRCC4-SIRF PLA foci in control (sgHPRT1) and ABRAXAS KO HT-29 cells treated with the indicated doses of CPT+/- ATMi. Data represent mean ± SEM from n ≥ 152 nuclei pooled from two independent experiments; *p* values are indicated, unpaired two-tailed t test. **e,** Representative images of metaphase chromosomes prepared from control (sgHPRT1) and ABRAXAS KO cells treated with CPT (2.5 nM) +/- ATMi (50 nM) for 16 h followed by 6 h colcemid treatment. **f,** Bar graph showing quantification of chromosomal abnormalities in cells as described in (e). Data represent mean ± SEM of n=3 independent experiments. At least 60 metaphase spreads were analyzed for each condition across three repeats; *p* values are indicated; unpaired two-tailed t test. **g,** Model illustrating how BRCA1-A complex restricts end resection by preventing fork reversal in ATM-inhibited cells. ATM inhibition results in compacted chromatin which favors BRCA1-A complex engagement at damaged replication forks. BRCA1-A complex prevents fork reversal resulting in eventual replication collapse. Illegitimate engagement of NHEJ at collapsed forks leads to heightened genomic instability and Top1i hypersensitivity (left). BRCA1-A complex loss in ATM-inhibited cells promotes chromatin accessibility at damaged forks and restores fork reversal catalyzed by FBH1, HLTF and ZRANB3 to generate reversed fork substrates. Controlled resection of a reversed fork generates 3’-ssDNA that can invades parental duplex ahead of the fork by HDR mechanism^49, 68, 76^. This leads to fork rescue, resulting in cell viability and Top1i resistance (right).

Impaired fork reversal results in fork breakage due to replication runoff at Top1 cleavage complexes (Top1cc)^46^. NHEJ-mediated illegitimate joining of broken DNA ends leads to chromosomal aberrations, radial chromosomes, and cytotoxicity^44, 77^. Because BRCA1-A prevents fork reversal upon ATMi (**Fig. 5b)**, we determined whether this leads to increased fork engagement of NHEJ factors. ATMi increased XRCC4-EdU PLA signal at both low (50 nM) and high (1 µM) CPT dose (**Fig. 7d**). However, ABRAXAS KO suppressed XRCC4 fork abundance in ATM-inhibited cells. Consistent with previous reports in *ATM*-null mouse cells, metaphase chromosome analysis further revealed a striking increase in chromosomal aberrations in ATMi treated cells, with a high abundance of radial structures (**Fig. 7e, 7f**). These were significantly reduced following ABRAXAS KO (**Fig. 7e, 7f**). ATMi-induced chromosomal aberrations were also found to be reduced in cells expressing RAP80 lacking both SIM and UIM domains (**Extended Data Fig. 12c)**. These results are consistent with a model that BRCA1-A recognition of damaged forks in ATMi treated cells drives heightened chromosomal aberrations via toxic NHEJ (**Fig. 7g**).

## Discussion

ATM kinase activity is central to DNA repair and required for cell survival following replication damage^39, 78–81^. Accordingly, pharmacological ATM inhibition produces cellular hypersensitivity to fork damaging agents, Top1i or PARPi, and these combinations are currently being investigated in clinical trials (NCT02588105). In this study, we show that BRCA1-A prevents fork reversal in ATM-deficient cells and restricts the generation of end resection substrates required for damaged fork repair. Our results support a model where ATMi promotes BRCA1-A engagement at damaged forks via SUMO and ubiquitin recognition, resulting in fork reversal blockade and eventual fork collapse (**Fig. 7g**). This engages toxic NHEJ at collapsed forks leading to chromosomal aberrations and cell death. Conversely, BRCA1-A loss restores fork reversal catalyzed by FBH1, HLTF or ZRANB3 ATP-dependent translocases, thereby generating substrates for controlled nucleolytic processing and subsequent fork recovery by homology-directed repair (HDR).

Loss of either NHEJ factors or BRCA1-A suppresses drug hypersensitivity in ATM-deficient cells, suggesting a critical role of BRCA1-A complex in governing repair outcome following replication fork damage in these genetic settings^35, 38, 39, 43, 82–84^. This represents a mechanistically distinct resistance process from BRCA1-deficient cancers where 53BP1 loss confers chemoresistance^44, 45^. It is noteworthy that BRCA1 plays dichotomous roles in each genetic context, highlighting its versatility in the DNA damage response through differential recognition of chromatin modifications. BRCA1-A prevents resection dependent damaged fork recovery in the context of ATM deficiency whereas 53BP1 loss is required to restore end resection and HR in BRCA1 mutant cells to cause PARPi or Top1i resistance. These differences further suggest that BRCA1-A and 53BP1 recognize different structures at damaged chromatin to suppress end-resection. 53BP1 requires RNF168 dependent H2AK15Ub and histone H4K20Me2 for damage recognition and suppression of end resection by nucleating the Shieldin complex. BRCA1-A localization to damaged forks required dual SUMO-ubiquitin recognition coincident with increased resection and CPT resistance in RAP80 SIM+UIM deleted mutants. BRCA1 interaction with ABRAXAS was also necessary for fork recognition and hypersensitivity to CPT+ ATMi. This implicates BRCA1-BARD1 damage recognition elements in association with the A-complex in creating the condensed chromatin structure that suppresses fork reversal and generation of end resection substrates. The BARD1-BRCT interacts with H2AK15Ub, and its Ankyrin repeats bind H4K20Me0 that are deposited on newly replicated chromatin^29, 30^. Along with RAP80 binding to SUMO-ubiquitin, BARD1 recognition of unmethylated histone H4 on nascent chromatin may further contribute to the different role of BRCA1-A in comparison to 53BP1 in limiting resection at damaged replication forks.

Our findings provide insights into how ATM regulates chromatin state of replication forks in response to DNA damage. iPOND-MS analysis revealed increased abundance of resection nucleases along with chromatin remodeling factors upon BRCA1-A loss in ATM-inhibited cells. This argues that BRCA1-A prevents fork resection by restricting nuclease access to damaged forks via regulating chromatin accessibility. This notion was buttressed by increased chromatin accessibility coupled with reduced H3K9me3 marks at damaged forks upon BRCA1-A loss. ATM signaling promotes chromatin accessibility surrounding DSB sites, particularly those located within heterochromatin^62, 63, 85^. On the other hand, recent findings suggest that replication stress induces a closed chromatin conformation that facilitates fork recruitment of BRCA1-BARD1 and inhibits nucleolytic processing^64^. We envision that chromatin compaction upon ATMi favors BRCA1-A engagement at damaged forks in a manner dependent on increased chromatin SUMOylation and ubiquitination. This in turn creates a stable chromatin state aided by multivalent nucleosome interactions and nucleosome bridging mediated by the BRCA1-BARD1 heterodimer in association with the A-Complex^31, 86^. Collectively, this enforces a block to fork reversal and access of end resection nucleases, as fork reversal involves major structural remodeling that inevitably requires changes to the local chromatin state of stressed replication forks ^68, 87^. The enhanced end resection and resistance to combined CPT and ATMi treatment in BRCA1-A deficient cells were alleviated upon genetic deletion of MRE11 and EXO1 nucleases. This agrees with recent findings demonstrating reliance of BRCA1 but not BRCA2 mutated cancer cells on EXO1 nuclease for survival^88, 89^. Although mechanistically distinct, our results further establish restored end resection as a key determinant of viability in both BRCA1 and ATM mutated cancers^39, 44^.

ATM plays a poorly understood role in promoting end resection at damaged forks^43, 83, 90^. Our study demonstrated a direct role of ATM kinase in generating resection substrates by promoting fork reversal following replication damage. Using electron microscopy, we provide direct experimental evidence of how damaged replication fork reversal is modulated by the ATM and BRCA1-A status. ATMi markedly reduced fork reversal, whereas BRCA1-A loss substantially restored it in ATM inhibited cells. Thus, BRCA1-A loss generates reversed fork substrates for resection in ATM-deficient cells. These observations present non-mutually exclusive possibilities that ATM directs fork reversal via either removal of BRCA1-A by direct phosphorylation or through phosphorylation of other substrates that affect fork SUMOylation, ubiquitylation and chromatin compaction ^13, 19, 20, 55, 91–96^. While the fork protection function of BRCA1 is well established, it does not appear to affect fork reversal^97–99^. Our study highlights a unique function of BRCA1-A in ATM-deficient background in blocking fork reversal, thereby suppressing end resection. This also highlights how generation of resection substrates following replication stress are regulated in a manner that depends on different genetic contexts^100^.

Top1i induces replication fork reversal, a RAD51-catalyzed intrinsic protective mechanism that prevents fork collapse into one-ended DSBs following replication stress^46, 47, 101, 102^. In addition to RAD51, fork reversal typically requires ATP-dependent DNA translocases including SMARCAL1, ZRANB3, HLTF or FBH1^69, 100, 103–106^. Nascent DNA ends of the reversed forks structurally resemble one-ended DSBs and engage nucleases for nucleolytic processing. Diminished ssDNA generation upon genetic perturbation of FBH1, HLTF or ZRANB3 translocases or RAD51 recombinase in BRCA1-A deficient cells supports the argument that fork reversal generates resection substrates upon BRCA1-A loss. Replication stalling can also result in ssDNA gaps generated by PRIMPOL mediated repriming downstream of DNA lesions^71–73^. These gaps are vulnerable for MRE11 and EXO1-mediated processing leading to bidirectional gap expansions^74, 75^. However, enhanced resection in BRCA1-A deficient cells did not require PRIMPOL. Together, these findings argue that BRCA1-A prevents end resection at reversed fork structures but not ssDNA gaps in ATM-deficient cells.

DNA end resection at stressed forks could be either physiological or pathological, depending on genetic context ^49, 51, 65, 99, 107, 108^. We show that nucleolytic processing of reversed forks upon BRCA1-A loss elicits a pro-survival response in ATM-deficient backgrounds. This agrees with published results demonstrating that controlled nucleolytic resection of reversed forks is indispensable for damage repair and replication restart^48, 49, 65^. Consequently, fork reversal is a critical determinant of genome stability in ATM deficient cells and may represent a therapeutic target to either reduce or exacerbate replication stress in ATM mutated individuals and associated cancers, respectively.

## Methods

### Cell lines

HT-29 and HEK293T cell lines were purchased from ATCC. Cell lines were cultured in 1X Dulbecco’s modified Eagle’s medium (GIBCO) supplemented with 10% bovine calf serum (Corning) and 1% Pen Strep (Gibco) at 37^°^C with 5% CO_2_. SMARCAL1, ZRANB3, HLTF and FBH1 KO U2OS cell lines were kindly provided by David Cortez, Vanderbilt University School of Medicine.

### CRISPR-Cas9 gene editing

CRISPR-based gene knockout was carried out using either *Streptococcus pyogenes* Cas9 (SpCas9) or *Staphylococcus aureus* Cas9 (SaCas9) system. For SpCas9, stable cell lines expressing SpCas9 were first generated by lentiviral transduction with lenti-SpCas9 hygro (Addgene, plasmid #104995), and subsequent selection with hygromycin (500 µg/ml). sgRNA guides were cloned into either lentiGuide-Puro (Addgene, plasmid #52963) or lentiGuide-neo^109^ (U6-sgRNA-P2A-Neo, reconstructed from Addgene plasmid #52963) vector digested with BsmBI-v2 (New England Biolabs) following the protocol described previously^110^. sgRNA plasmids (2.7 µg) were packaged into lentivirus particles by co-transfecting with packaging plasmids psPAX2 (2 µg, Addgene, plasmid #12260) and pMD2.G (1.3 µg, Addgene, plasmid #12259) into HEK293T cells grown in a 6-well plate using lipofectamine 2000 (Invitrogen). Lentivirus supernatant was collected 48 h post-transfection, filtered through a 0.45-µm filter (Millipore) and subsequently used to transduce the target cells using polybrene (8 µg/ml, Sigma-Aldrich) by spin infection at 2000 rpm for 90 min. Stable transduced cells were selected by either puromycin (4 µg/ml) or neomycin (800 µg/ml), and knockout efficiencies were verified by Western blotting. ABRAXAS KO HEK293T cells were generated by transducing spCas9-expressing (lenti-SpCas9 hygro) stable HEK293T cells with lentivirus particles carrying ABRAXAS sgRNA guides cloned into lentiGuide-Puro plasmid. ABRAXAS KO HT-29 cells were generated by infecting spCas9-expressing stable HT-29 cells with lentivirus particles harboring ABRAXAS sgRNA guides cloned into lentiGuide-neo vector. For generating ABRAXAS KO HT-29 cells using the SaCas9 system, cells were transduced with lentivirus particles carrying ABRAXAS sgRNA guides cloned into SaLgCn (U6-sgRNA-EFS-SaCas9-P2A-Neo^77^) plasmid engineered to express both SaCas9 and sgRNA. Control sgROSA and sgHPRT1 harboring cell lines were generated using SpCas9 and SaCas9 systems, respectively. To make ATM-ABRAXAS, MRE11-ABRAXAS, EXO1-ABRAXAS and PRIMPOL-ABRAXAS double KO cells, ABRAXAS was knocked out using SaCas9, while SpCas9 system was used to KO the respective genes. ABRAXAS was KO in SMARCAL1, ZRANB3, HLTF or FBH1 KO U2OS cells using SaCas9 system^100^. For MRE11 knockout, experiments were conducted 48 h following puromycin selection. sgRNA guide sequences are provided in Supplementary Table 1.

### Site-directed mutagenesis and generation of stable cell lines

ABRAXAS and RAP80 mutants were generated by site-directed mutagenesis of pOZ-N-FH Abraxas^111^ (Addgene, Plasmid #27495) and pOZ-N-FH RAP80^111^ (Addgene, Plasmid #27501) retrovirus expression plasmids, respectively using Q5 Site-Directed Mutagenesis kit (New England Biolabs, E0554S). Retrovirus particles were generated by co-transfection of the pOZ-expression plasmids and pCL-Ampho packaging vector with 2:1 ratio into Phoenix cells using lipofectamine 2000 (Invitrogen). Retrovirus supernatant was collected 48 h post-transfection and filtered through a 0.45 µm filter. ABRAXAS or RAP80 KO cells were transduced with pOZ-retrovirus particles using polybrene (8 µg/ml, Sigma-Aldrich) by spin infection at 2000 rpm for 90 min. Stably transduced cells were selected on anti-IL2-receptor antibody coated magnetic beads 48 h after virus infection and overexpression was confirmed by Western blotting.

### siRNA transfection

HT-29 cells were transfected two consecutive days with either non-targeting control siRNA (D-001810-01-05, ON-TARGETplus, Horizon Discovery) or siRNA targeting RAD51 (L-003530-00-0005, Horizon Discovery) at a final concentration of 100 nM using DharmaFECT 1 transfection reagent (Horizon Discovery). Cells were fixed for immunofluorescence experiments 72 h after the 1^st^ transfection and RAD51 knockdown was verified by Western blotting.

### iPOND mass spectrometry

iPOND was performed as described previously with modifications^112^. Control (sgROSA) and ABRAXAS KO HEK293T cells (∼100 X 10^6^ cells for each genotype) treated with CPT plus ATMi were pulse labeled with 10 µM EdU for 20 min. An additional set of control cells were kept for no-click negative control. Cells were fixed immediately with 1% formaldehyde in 1X PBS for 20 min at RT followed by glycine (1.25 M) quenching. Cells were collected by scraping and pelleted by centrifugation at 900 x g for 5 min at 4°C. Cells were washed three times with 1X ice-cold PBS followed by permeabilization with 0.25% Triton X-100 in PBS for 30 min at RT. Cells were pelleted by centrifugation at 900 x g for 5 min at 4 °C and washed with 0.5% BSA in 1X PBS. Cell pellets were resuspended in click reaction cocktails (20 μM biotin-azide, 2 mM CuSO4, 10 mM sodium L-ascorbate in 1X PBS), vortexed and incubated for 2 h at RT with rotation. For no-click negative control, biotin-azide was not included in the click reaction cocktails. Cells were pelleted by centrifugation at 900 x g for 5 min at 4 °C and washed with 0.5% BSA in 1X PBS. Cell pellets were resuspended in lysis buffer (1% SDS in 50 mM Tris-HCl, pH 8.0) at a concentration of 100 X 10^6^ cells per 4 ml and sonicated using a microtip sonicator (Branson 150) set at 35% amplitude with 5 pulses (20 S on and 40 S pause). Lysates were clarified twice by centrifugation at 16100 x g for 10 min at RT, filtered through an 80 µm nylon membrane (NY8004700, Millipore Sigma) and diluted 1:1 (v/v) with 1X ice-cold PBS containing protease inhibitors. Lysates were incubated with streptavidin agarose beads (69203, Millipore Sigma) at a concentration of 50 µl (packed volume) beads per 100 X 10^6^ cells overnight at RT with rotation. Beads were washed once with cold lysis buffer, once with 1 M NaCl and final two washes with cold lysis buffer. Proteins were eluted by boiling the beads in 4X LDS sample buffer (NP0007, Thermo Scientific) at 95°C, and iPOND purification was evaluated by either silver staining or Western blotting using antibodies for replication fork associated proteins.

Mass spectrometry analysis of iPOND purified protein samples was carried out in Taplin Biological Mass Spectrometry Facility, Harvard Medical School. Briefly, beads were washed at least five times with 100 µl 50 mM ammonium bicarbonate then 5 µl (200 ng/µl) of modified sequencing-grade trypsin (Promega, Madison, WI) was spiked in and the samples were placed in a 37°C room overnight. The samples were then centrifuged and the liquid removed. The extracts were then dried in a speed-vac (∼1 h). Samples were re-suspended in 50 µl of HPLC solvent A (2.5% acetonitrile, 0.1% formic acid) and desalted by STAGE tip^113^. On the day of analysis, the samples were reconstituted in 10 µl of HPLC solvent A. A nano-scale reverse-phase HPLC capillary column was created by packing 2.6 µm C18 spherical silica beads into a fused silica capillary (100 µm inner diameter x ∼30 cm length) with a flame-drawn tip^114^. After equilibrating the column, each sample was loaded via a Famos auto sampler (LC Packings, San Francisco CA) onto the column. A gradient was formed, and peptides were eluted with increasing concentrations of solvent B (97.5% acetonitrile, 0.1% formic acid). As peptides eluted, they were subjected to electrospray ionization and then entered into a Velos Orbitrap Elite ion-trap mass spectrometer (Thermo Fisher Scientific, Waltham, MA). Peptides were detected, isolated, and fragmented to produce a tandem mass spectrum of specific fragment ions for each peptide. Peptide sequences (and hence protein identity) were determined by matching protein databases with the acquired fragmentation pattern by the software program, Sequest (Thermo Fisher Scientific, Waltham, MA)^115^. All databases include a reversed version of all the sequences and the data was filtered to between a one and two percent peptide false discovery rate. For fold change calculation, normalization was done by multiplying each spectrum count in a sample by the average number of spectra across all samples/total number of spectra in the respective sample. Proteins with ≥ 6 total spectrum count detected in control and ABRAXAS KO cells across all replicates, and showed > 5 average fold change in iPOND samples over no click (NC) negative control were considered for fold change analysis. Log2 fold change was calculated from *n*=5 biological replicates and scatter plot was generated in GraphPad Prism 10.

### Gene ontology (GO) analysis

Gene ontology analysis of proteins identified from iPOND-MS analysis was done using GeneCodis 4^116^. Proteins enriched by ≥ 1.20-fold in ABRAXAS KO cells over control (sgROSA) cells were analyzed for GO biological processes and GO molecular functions. GO enrichment analysis data are provided in Supplementary Table 3.

### Colony survival assay

Cells were plated (2000 cells) on 6-well tissue culture plates and treated with CPT and ATMi 4 h after cell seeding. Cells were allowed to form colonies for two to three weeks. After 1X PBS wash, colonies were fixed with 100% methanol, subsequently stained with staining solution (25% methanol+ 0.5% crystal violet). The total number of colonies was normalized with plating efficiency and relative colony survival was determined as percentage of colonies survived following drug treatments. Graphs were generated in MS Excel.

### Immunofluorescence

Cells grown onto coverslips or 8-well chamber slides (229168, Celltreat) were treated with CPT and ATMi as indicated in the figures. Cells were washed twice with ice-cold 1X PBS followed by nuclear pre-extraction for 10 min on ice using CSK buffer (10 mM PIPES, pH 7.0; 100 mM NaCl; 300 mM Sucrose; 3 mM MgCl_2_; 1 mM EGTA, 0.5% Triton X-100). Cells were washed twice with 1X PBS and fixed with 4% paraformaldehyde (Electron Microscopy Sciences, 50-980-495) diluted in 1X PBS solution for 20 min on ice. Cells were washed three times with ice-cold 1X PBS, permeabilized with 0.5% Triton X-100 for 10 min on ice and incubated in blocking buffer (5% BSA with 10% goat serum in 1X PBS) at 4°C for 30 min. Cells were incubated in primary antibodies diluted in blocking buffer overnight at 4°C. Cells were washed three times with 1X PBS followed by incubation in AlexaFluor 647- or AlexaFluor 488-conjugated secondary antibodies (Thermo Scientific) for 1 h at RT. Cells were washed three times with 1X PBS, stained with DAPI (D9542, Sigma-Aldrich, 4 µg/ml diluted in 1X PBS) and mounted with ProLong Gold Antifade mountant (P36930, Thermo Scientific). Immunofluorescence images were acquired by CoolSnap MYO camera of Nikon Eclipse 80i fluorescence microscope using a Plan Apo VC 60× Oil DIC N2 objective lens. Images with 100X magnification were acquired using a 100X oil objective lens of Nikon Eclipse Ti2 microscope. Images were quantified using CellProfiler 4.2.5 image analysis software^117^ and graphs were generated in GraphPad Prism 10. Antibodies used in immunofluorescence experiments are listed in Supplementary Table 4.

### BrdU-coupled MNase accessibility assay

HEK293T cells were treated with CPT (1 µM) and ATMi (500 nM) and pulse labeled with 20 µM BrdU for 12 min as indicated in Figure 2A. Cells were washed with 1X ice-cold PBS, trypsinized and pelleted by centrifugation at 300g for 10 min at 4 °C. Equal number of cells (∼30 X10^6^) in each sample were resuspended in 5 ml of ice-cold NP-40 lysis buffer (10 mM TrisHCl pH 7.4, 10 mM NaCl, 3 mM MgCl2, 0.5% Nonidet P-40, 0.15 mM spermine, and 0.5 mM spermidine), incubated on ice for 5 min and centrifuged at 120g for 10 min at 4 °C. The nuclei were washed with 2.5 ml of MNase digestion buffer (10 mM Tris-HCl (pH 7.4), 15 mM NaCl, 60 mM KCl, 0.15 mM spermine and 0.5 mM spermidine) and pelleted by centrifugation for 10 min 4 °C. The nuclei were resuspended in 1 ml of MNase digestion buffer containing 1 mM CaCl2, and 100 µl of resuspended nuclei were subjected to MNase (250 gel units) digestion at 37 °C for the time periods indicated in Figure 2. Reactions were stopped by adding MNase stop buffer (100 mM EDTA and 10 mM EGTA pH 7.5) diluted 1:5 in digestion buffer. Samples were incubated at 37 °C overnight after the addition of 3 µl of proteinase K (25 mg/ml., Invitrogen, EO0491) and 10 µl of 20% SDS. Genomic DNA was purified by Phenol chloroform extraction methods and ethanol precipitation. Equal amount of DNA (4 µg) in each sample was run on 1.2% agarose gel, stained with EtBr and photographed using a GelDoc system (Biorad). The gel was treated with 0.2 N HCL for 10 min, washed several times in water and treated with denaturation solution (1.5 M NaCl, 0.5 M NaOH) for 30 min with gentle rocking. The gel was neutralized by neutralization solution (3 M NaCl, 0.5 M Tris pH 7.5) for 30 min and subjected to Southern transfer to Hybond-N+ charged nylon membrane (RPN203B, Cytiva) using 20X SSC buffer (3M NaCl, 0.3 M sodium citrate) at RT overnight. Membrane was crosslinked in a UVP crosslinker (120 mj/cm^2^), blocked in 5% milk for 1 h and incubated in anti-BrdU antibody (1:200, 347580, BD Biosciences) overnight at 4 °C. Membrane was washed three times with 1X PBST (Phosphate Buffer Saline with 0.1% Tween 20), incubated in anti-mouse secondary antibody for 1h at RT and washed 5 times with 1X PBST. BrdU labeled DNA fragments were detected using Western lightning plus-ECL (NEL105001EA, Revvity Health Sciences Inc.). BrdU labeled nucleosomes were quantified in ImageJ (https://imagej.net/ij/) and MNase accessibility was determined by fractions of BrdU labeled mononulceosomes as a percentage of total BrdU-labeled DNA in each lane. Graphs were generated in GraphPad Prism 10

### SIRF assay

SIRF assay was performed as described previously with modifications^118, 119^. Control (sgHPRT1) or ABRAXAS KO HT-29 cells were plated in 8-well chamber slides at 60% confluency the day before the experiments. Cells were optionally pretreated with 250 nM ATMi for 10 min followed by incubation in CPT (1 µM or 50 nM) containing medium for 40 min. Cells were pulse labeled with 125 µM of EdU (A10044, Invitrogen) 20 min prior to the end of CPT treatment. ATM inhibitor was present during the entire experiment. Cells were washed two times with ice-cold 1X PBS followed by pre-extraction with CSK buffer (10 mM PIPES, pH 7.0; 100 mM NaCl; 300 mM Sucrose; 3 mM MgCl_2_; 1 mM EGTA, 0.5% Triton X-100) for 10 minutes on ice. Cells were fixed with 4% paraformaldehyde (Electron Microscopy Sciences, 50-980-495) for 20 min on ice. Cells were washed two times with ice-cold 1X PBS and permeabilized with 0.25% Triton X-100 for 10 min on ice. After washing two times with 1X PBS, cells were subjected to click reaction using 20 μM biotin-azide, 2 mM CuSO4, 100 mM sodium L-ascorbate diluted in 1X PBS for 1 h at RT. After washing with 1X PBS, slides were blocked with blocking buffer (5% BSA with 10% goat serum in 1X PBS) for 1 h at 4 °C followed by primary antibody incubation at 4 °C overnight. For MRE11 SIRF, fixed cells were incubated with mouse anti-MRE11 (GTX70212, GeneTex) and rabbit anti-biotin (5597, Cell Signaling) primary antibodies, whereas EXO1 SIRF was carried out using rabbit anti-EXO1 (GTX109891, GeneTex) and mouse anti-biotin antibodies (SAB4200680, Sigma-Aldrich). SUMO2/3-EdU and FK2 (Ub)-EdU SIRF experiments were carried out using mouse anti-Sumo 2/3 (ab81371, Abcam) and mouse anti-FK2(Ub) (04-263, Millipore Sigma) antibodies respectively in combination with rabbit anti-biotin (5597, Cell Signaling). An EdU-EdU SIRF was included to determine the total EdU incorporation in each experimental condition using mouse anti-biotin and rabbit anti-biotin antibodies. Slides were washed three times with wash buffer A (0.01 M Tris, 0.15 M NaCl, and 0.05% Tween 20, pH 7.4) followed by incubation with anti–mouse plus and anti–rabbit minus PLA probes diluted 1:5 in blocking solution for 1 h at 37°C. After three washes in wash buffer A, ligation reaction was performed at 37°C for 30 min using Duolink ligation buffer (diluted 1:5 in water) and ligase (diluted 1:40 in water) enzymes provided with Duolink® In Situ Detection Reagents Red kit (DUO92008, Sigma-Aldrich). Slides were briefly washed two times with wash buffer A and subjected to rolling circle amplification at 37°C for 100 min using amplification mix containing Duolink amplification reagent (diluted 1:5 in water) and rolling circle polymerase (diluted 1:80 in water). Slides were washed three times with wash buffer B (0.2 M Tris and 0.1 M NaCl) and one time with 0.01× diluted wash buffer B. Slides were stained with DAPI (D9542, Sigma Aldrich, 4 µg/ml diluted in 1X PBS) and mounted with ProLong Gold Antifade mountant (P36930, Thermo Scientific). Images were acquired by Nikon Eclipse Ti2 fluorescence microscope using a 100× Oil objective lens or a 60× Oil DIC N2 objective lens of Nikon Eclipse 80i fluorescence microscope. SIRF-PLA foci were quantified using CellProfiler 4.2.5 software and normalized with the average EdU-EdU signal in each sample. Scatter plots were generated in GraphPad Prism 10.

### Electron microscopy analysis

EM analysis of replication intermediates was performed as described previously^71, 120, 121^. Briefly, control and ABRAXAS KO HT-29 cells (∼60 X10^6^ cells in each sample) were treated with 50 nM CPT+/- 250 nM ATMi as shown in Figure 6A. Cells were washed with 1X ice cold PBS, collected by trypsinization and pelleted by centrifugation at 2000 rpm for 5 min at 4 °C. Cells were resuspended in 1X ice-cold PBS, transferred to 60 mm tissue culture dishes placed on a cold metal block. Genomic DNA was psoralene-crosslinked by addition of 4,5′,8-Trimethylpsoralen (TMP) in a spread-out fashion at a final concentration of 10 µg/ml and irradiated in a UVP crosslinker at 120 mj/cm^2^ for 3 min. The psoralene-crosslinking was repeated two more times and cells were pelleted by centrifugation at 2000 rpm for 5 min at 4 °C. Cells were washed three times with 1X ice-cold PBS, resuspended in cold lysis buffer (1.28 M Sucrose, 40 mM Tris-HCl pH 7.5, 20 mM MgCl2, 4% Triton X-100; diluted 1:4 with water) in a 1:4 ratio and incubated on ice for 10 min. Lysed cells were pelleted by centrifugation at 1500 rpm for 15min at 4 °C. After washing once with cold lysis buffer, nuclei were resuspended in 200 µl cold 1X PBS and subsequently resuspended in 5 ml of digestion buffer (800 mM guanidine-HCl, 30 mM Tris-HCl pH 8.0, 30 mM EDTA pH 8.0, 5% Tween-20, 0.5% Triton X-100). Samples were subjected to Proteinase K digestion (200 µl of 20 mg/ml) at 50 °C for 2 h. Genomic DNA was extracted by standard chloroform:isoamylalcohol and isopropanol precipitation method and digested with PvuII HF (R3151S, New England Biolabs) restriction enzyme at 37°C for 4h. Replication intermediates were enriched on plasmid miniprep columns (12123, QIAGEN) and subsequently concentrated using Amicon Ultra size exclusion columns (UFC510096, Millipore Sigma). For EM visualization, samples were prepared by spreading the concentrated DNA on a carbon-coated grid in presence of benzyl-dimethyl-alkylammonium chloride (B6295, Millipore Sigma) as well as n,n-dimethylformamide (227056, Millipore Sigma), followed by low angle platinum rotary shadowing. Images were acquired with a JEOL JEM-1400 electron microscope using a bottom mounted AMT XR401 camera. Image analysis was done using ImageJ software^122^. EM analysis allows distinguishing duplex DNA from ssDNA. DNA duplex is expected to appear as a 10 nm thick fiber following platinum coating step necessary for EM visualization, whereas ssDNA has a reduced thickness of 5-7 nm and appears lighter in contrast to that of dsDNA. Criteria used for the assignment of a three-way junction, indicative of a typical replication fork, include the joining of three DNA fibers into a single junction, with two generally symmetrical daughter strands and a single parental strand. Reversed replication forks comprise four DNA strands joined at a single junction. This consists of two commonly symmetrical daughter strands, one parental strand and an additional typically shorter fourth strand, representative of the reversed arm. The length of the two daughter strands corresponding to the newly replicated duplex oftentimes are equal (b=c), whereas the length of the parental arm and the regressed arm can vary (a ≠ b = c ≠ d). Conversely, canonical Holliday junction structures are characterized by arms of equal length (a = b, c = d). Particular attention was given to the junction of the reversed replication fork to observe the presence of a bubble structure, indicating that the junction is opened and that it was simply not the result of the occasional crossover of two DNA molecules. These four-way junctions of reversed replication forks may also be collapsed and other indicators such as daughter strand symmetry, presence of single-stranded DNA at the junction or the entire structure itself, all are considered during analysis^121^. The frequency of reversed forks, junction gaps and daughter strand gaps in a sample was computed using Prism software.

### Detection of nascent ssDNA by IdU under non-denaturing conditions

To detect nascent ssDNA, cells were treated with 50 nM CPT +/- 250 nM ATMi and concomitantly pulse labeled with 250 µM IdU. Nuclei were pre-extracted, fixed and permeabilized as described under Immunofluorescence method section. After incubation in blocking buffer (5% BSA with 10% goat serum in 1X PBS) for 30 min at 4 °C, fixed cells were incubated with mouse anti-IdU (1:100, 347580, BD Pharmigen) and rabbit anti-γH2AX (1:500, 39117, Active Motif) antibodies diluted in blocking buffer overnight at 4°C. After three PBS wash, cells were incubated with anti-mouse AlexaFluor 488 (1:500, A-11029, Thermo Scientific) and anti-rabbit AlexaFluor 647 (1:500, A-21245, Thermo Scientific) conjugated secondary antibodies for 1 h at RT. Cells were washed with 1X PBS, stained with DAPI (D9542, Sigma-Aldrich, 4 µg/ml diluted in 1X PBS) and mounted using ProLong Gold Antifade mountant (P36930, Thermo Scientific). IdU immunofluorescence foci were examined by Nikon Eclipse 80i fluorescence microscope using a 60× Oil DIC N2 objective lens. Images were quantified using CellProfiler 4.2.5 software and scatter plots were generated in GraphPad Prism 10.

### DNA fiber assay

Exponentially growing control (sgHPRT1) or ABRAXAS KO HT-29 cells (∼ 0.2 x 106) were sequentially labeled with 25 μM CldU (C6891, Millipore Sigma) and 250 μM IdU (I7125, Millipore Sigma), and concomitantly treated with 1 µM CPT +/- 250 nM ATMi as shown in Figure 7A. Cells were washed with ice-cold 1X PBS, trypsinized, and ∼2,000 cells were lysed using lysis buffer (200 mM Tris-HCl pH 7.4, 50 mM EDTA and 0.5% SDS) on a silane coated slides (5070, Newcomer Supply). DNA fibers were spread by gravity flow, air dried and fixed in fixative solution (Methanol: Acetic acid, 3:1). Slides were treated with 2.5 M HCl for 1 h at RT, neutralized with 400 mM Tris-HCl, pH 7.4. After PBST wash, slides were blocked with blocking buffer (5% BSA and 10% goat serum in 1X PBS) at 4°C overnight. Slides were incubated with rat anti-CIdU antibody (1:100, ab6326, Abcam) for 1 h at RT. After three PBST wash, slides were incubated with AlexaFluor 647-conjugated anti-rat secondary antibody (1:100, A-21247, ThermoFisher Scientific) for 1 h at RT and washed three times with PBST. Next, the slides were incubated with mouse anti-IdU antibody (1: 40, 347580, BD Pharmigen) for 1 h at RT, washed with PBST and subsequently incubated with AlexaFluor 488-conjugated anti-mouse secondary antibody (1:100, A-11029, ThermoFisher Scientific) for 1 h at RT. Slides were washed three times in PBST and mounted with Prolong Gold antifade mountant (P36930, ThermoFisher Scientific). DNA fibers were imaged by Nikon Eclipse 80i fluorescence microscope using a Plan Apo VC 60× Oil DIC N2 objective lens. Fibers were quantified in ImageJ (https://imagej.net/ij/) and fork restart efficiency was determined by fractions of CldU-IdU (red-green) labeled fibers as a percentage of total CldU (red and red-green) labeled fibers. ≥150 fibers were analyzed for each experimental condition. Bar graphs were generated in GraphPad Prism 10.

### Metaphase chromosome analysis

Metaphase chromosomes were prepared as described previously^77^. Briefly, HT-29 cells (5 × 10^6^) growing in 10 cm dishes were treated with 2.5 nM CPT in combination with 50 nM ATMi for 16 h followed by metaphase arrest using KaryoMAX Colcemid solution (0.1 μg/ml) (15212012, ThermoFisher Scientific) for 6 h. Cells were washed with ice-cold 1X PBS, trypsinized, and collected by centrifugation at 100×g for 5 min. Cell pellets were resuspended in 75 mM prewarmed (37 °C) KCL in a drop-wise manner using slow vortexing followed by incubation at 37 °C for 30 min. Cells were fixed two times by dropwise addition of 6 ml freshly prepared fixative solution (methanol: acetic acid, 3:1) with gentle vortexing. After centrifugation at 100×g for 5 min, cell pellets were resuspended in 2 ml fixative solution and stored at −20 °C overnight. Cells were resuspended in 1 ml freshly prepared fixative solution followed by metaphase chromosome preparation by dropping 200-300 μl of the cell suspension on silane-coated glass slide (5070, Newcomer Supply) presoaked in 40% acetic acid. The slides were air dried, recovered overnight and stained with Giemsa (48900, Sigma-Aldrich) followed by mounting using Prolong Gold antifade mountant (P36930, ThermoFisher). Images were acquired using Nikon Eclipse Ti2 fluorescence microscope equipped with a 100× Oil objective lens.

### Western blotting

Cells were lysed in 2X LDS sample buffer (NP0007, Thermo Scientific) supplemented with benzonase (E1014, Sigma-Aldrich) and 1.5 mM MgCl2. Proteins were separated onto NuPAGE™ 4-12% Bis-Tris (NP0335BOX, ThermoFisher Scientific) gels, transferred onto 0.2 µm nitrocellulose blotting membrane (10600004, Cytiva) by wet transfer at 100 constant volts for 2h at 4 °C. The membranes were blocked with 5% milk diluted in 1X PBS for 1 h followed by primary antibody incubation overnight at 4 °C. Membranes were washed three times with PBST and incubated with HRP-conjugated anti-mouse (NA931, Amersham, 1:2500 dilution) or anti-rabbit (NA934, Amersham, 1:2500 dilution) secondary antibodies at RT for 1 h. Membranes were washed with PBST, and protein bands were detected using Western lightning plus-ECL (NEL105001EA, Revvity Health Sciences Inc.). Antibodies used in Western blotting are listed in Supplementary Table 4.

## QUANTIFICATION AND STATISTICAL ANALYSIS

Unpaired two-tailed *t* test and two-tailed Mann–Whitney test were performed to compute statistical significance in GraphPad Prism 10. P values are indicated in the respective figures.

## Acknowledgments

We thank members of the Greenberg lab and Penn Center for Genome Integrity for critical discussion. This work was supported by NIH R01s 138835 and 174904, and funds from the Basser Center for BRCA to R.A.G., and R01CA237263 and R01CA248526 to A.V. We thank David Cortez, Department of Biochemistry, Vanderbilt University School of Medicine for the kind gift of SMARCAL1, ZRANB3, HLTF and FBH1 KO U2OS cells.

## Author contributions

A.D. performed the major experiments involving cellular hypersensitivity to DNA damaging agents, immunofluorescence, iPOND-mass spectrometry, DNA fiber analysis, and chromatin accessibility assays with assistance from J.Q. Y.I.M. performed initial camptothecin hypersensitivity assays using RAP80 knockout and reconstitution approaches. J.J. and A.V. performed all electron microscopy experiments and analysis. A.D. and R.A.G. wrote the manuscript with critical input from A.V. and J.J.

## Competing interests

The authors declare no competing interests.

## Extended Data Figures

**Extended Data Fig. 1:**
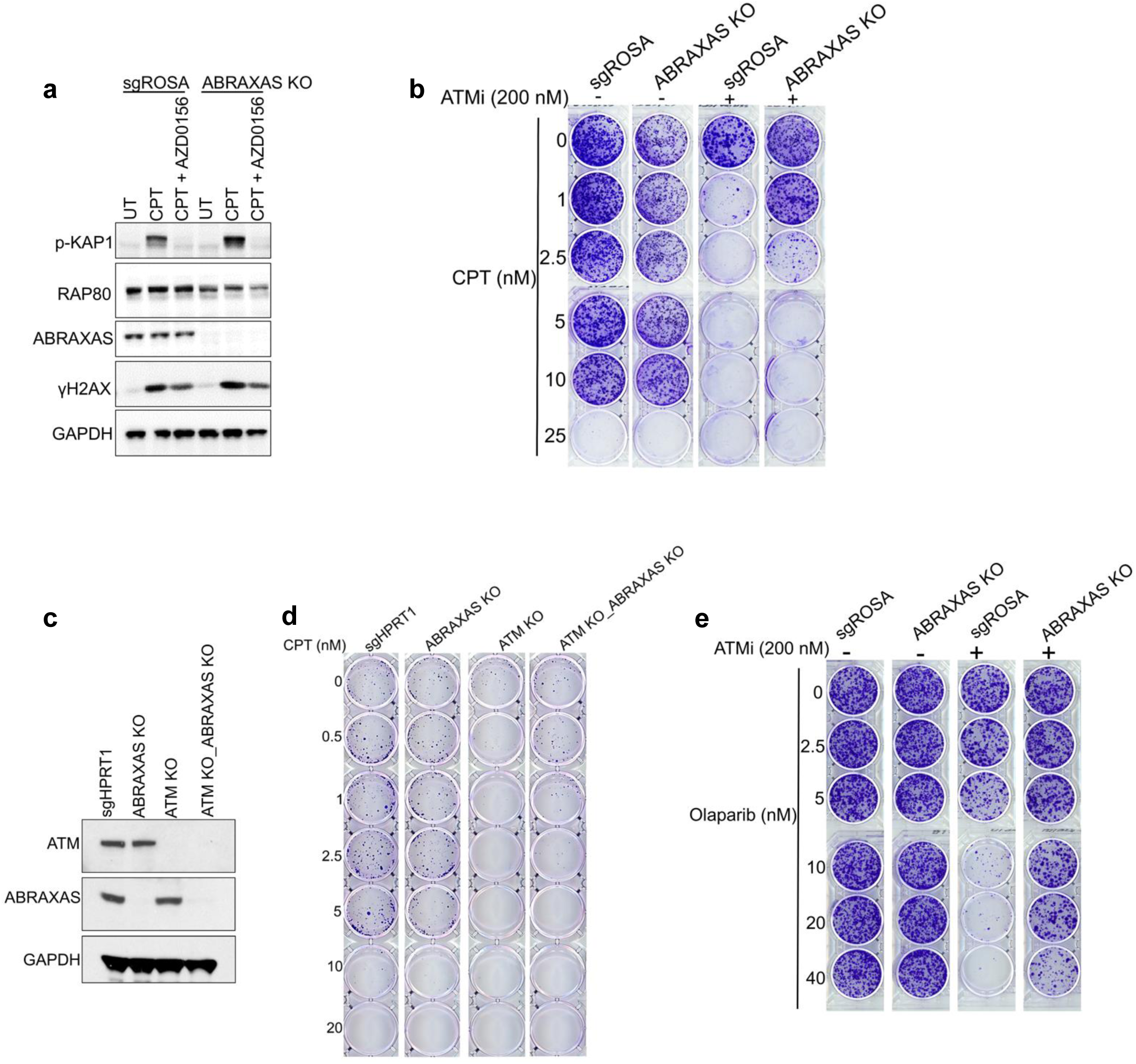
BRCA1-A complex loss confers drug resistance in ATM-deficient cancer cells. **a,** Western blotting showing diminished CPT-induced phospho (S824)-KAP1 (p-KAP1) levels upon AZD0156 treatment, confirming potency of the ATM inhibitor. Control (sgROSA) and ABRAXAS KO HEK293T cells were treated with CPT (1 µM) +/- AZD0156 (250 nM) for 40 min and analyzed by Western blotting using the indicated antibodies. GAPDH serves as loading control. **b,** Colony survival assay showing CPT+ATMi resistance upon ABRAXAS KO in HEK293T cells. Experiments were repeated *n*=3 times with similar results. **c,** Western blotting validating ATM and ABRAXAS DKO in HT-29 cells. GAPDH serves as loading control. **d,** Colony survival assay showing CPT resistance in ATM and ABRAXAS DKO HT-29 cells. Experiments were repeated *n*=3 times with similar results. **e,** Colony survival assay showing olaparib+ATMi resistance in ABRAXAS KO HEK293T cells. Experiments were repeated *n*=3 times with similar results.

**Extended Data Fig. 2:**
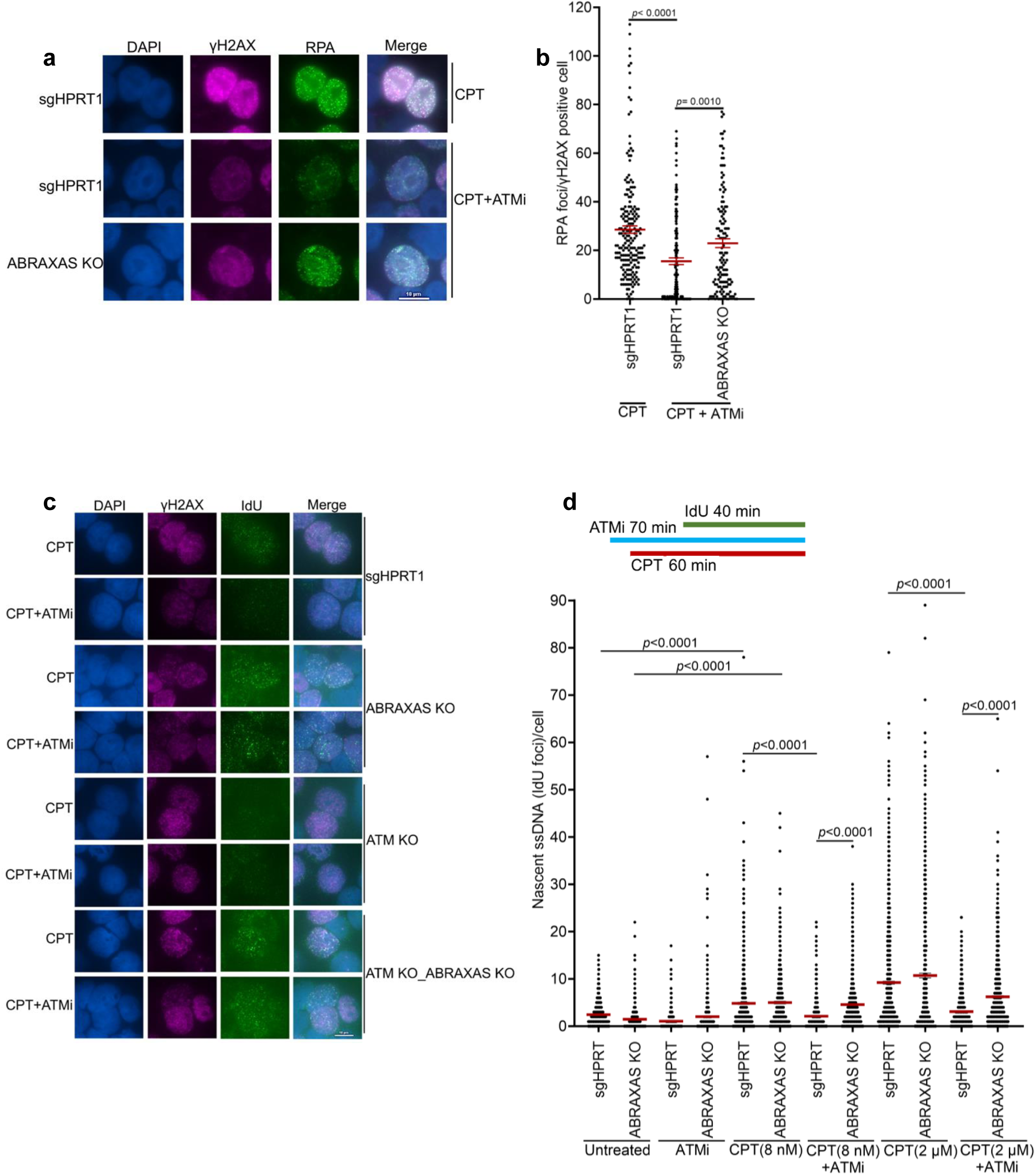
BRCA1-A complex suppresses damaged fork resection upon ATM inhibition. **(a and b),** Representative images of RPA immunofluorescence in control (sgHPRT1) and ABRAXAS KO HT-29 cells treated with CPT (1 µM) +/- ATMi (250 nM) for 1 h. ATMi was optionally added 10 min before CPT treatment and was present throughout the experiment. γH2AX immunostaining marks cells with CPT-induced replication damage. Scale bar is shown (**a**). Scatter plot showing quantification of RPA foci in γH2AX positive cells (**b**). Horizontal bars represent mean ± SEM derived from n ≥ 130 γH2AX positive cells examined over three independent experiments; *p* values are indicated, unpaired two-tailed t test. **c,** Representative images of anti-IdU immunofluorescence detected under non-denaturing conditions in the indicated genotypes of HT-29 cells treated with 50 nM CPT +/- 250 nM ATMi. Scale bar is shown. **d,** Schematic showing the experimental scheme of native IdU assay in control (sgHPRT1) and ABRAXAS KO HT-29 cells treated with low (8 nM) or high (2 µM) CPT +/- 250 nM ATMi. IdU was added 20 minutes after CPT addition to label newly synthesized DNA. Quantification of ssDNA (native IdU foci) is shown. Data represent mean ± SEM derived from n ≥ 390 cells examined over two independent experiments; *p* values are indicated, unpaired two-tailed t test.

**Extended Data Fig. 3:**
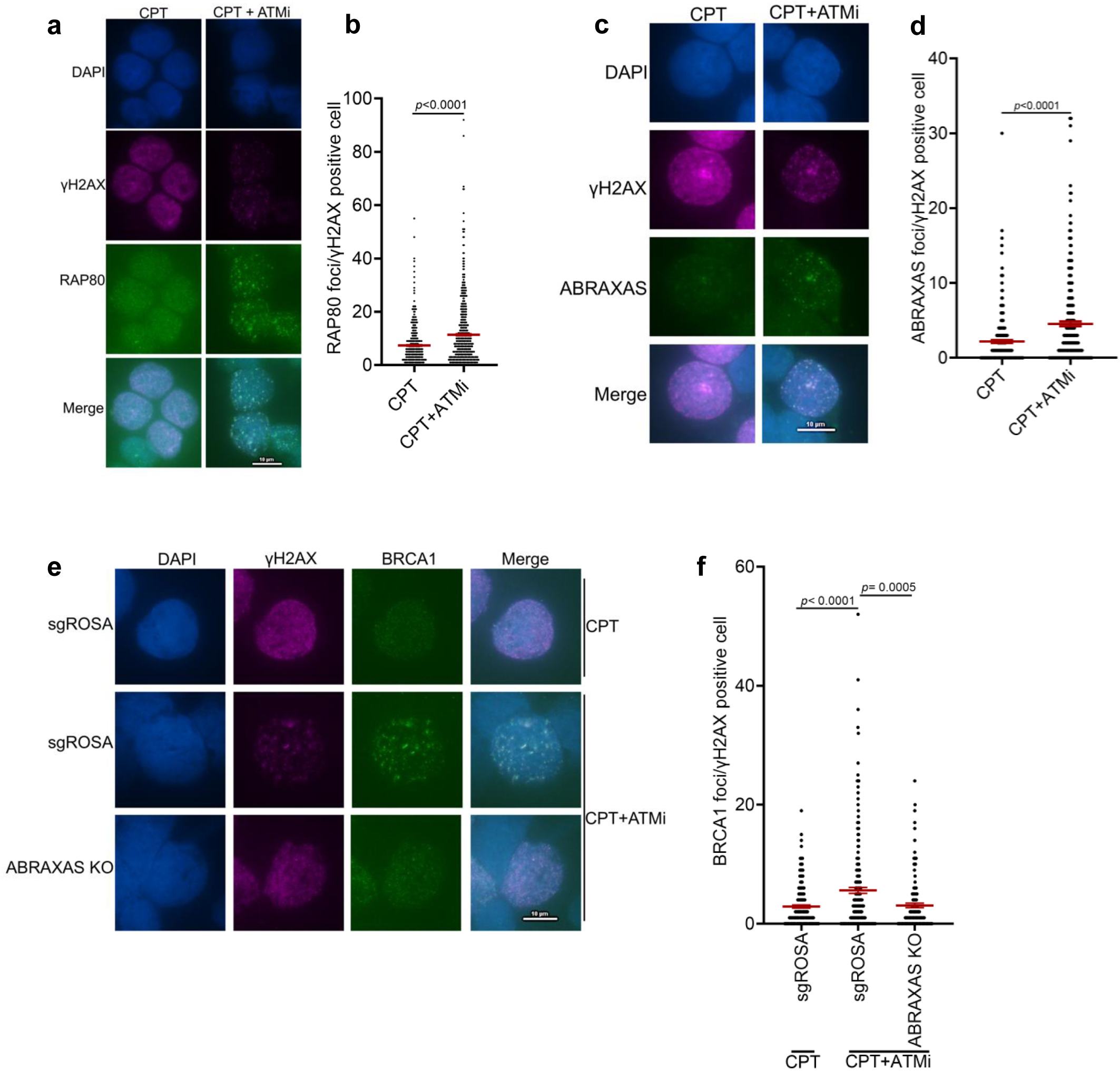
ATMi promotes BRCA1-A complex recruitment at damaged chromatin. **(a and b),** Representative immunofluorescence images of RAP80 foci in HT-29 cells treated with CPT (1 µM) alone or in combination with ATMi (250 nM) for 1 h. ATMi was added 10 min before CPT treatment (**a**). Quantification of the RAP80 foci is shown (**b**). Experiments were performed four times with similar results. Data represent mean ± SEM derived from n ≥ 325 cells examined over two independent experiments; *p* values are indicated, unpaired two-tailed t test. **c,** Representative immunofluorescence images showing increased HA-ABRAXAS foci in HT-29 cells upon ATM inhibition. Cells were treated with CPT (1 µM) +/- ATMi (250 nM) for 1 h. ATMi was added 10 min before CPT treatment. γH2AX marks cells with CPT-induced replication damage. Scale bar is shown. **d,** Quantification of the data from experiments as described in (c). Experiments were repeated three times with similar results. Data are mean ± SEM derived from n ≥ 300 cells examined over two independent experiments; p values are indicated, unpaired two-tailed t test. **(e** and **f),** Representative immunofluorescence images of BRCA1 foci in control (sgROSA) and ABRAXAS KO HT-29 cells treated with CPT (1 µM) +/- ATMi (250 nM) for 1 h. ATMi was added 10 min before CPT treatment (**e**). Scatter plot showing ABRAXAS-dependent increased BRCA1 foci in CPT-treated cells following ATM inhibition (**f**). Data represent mean ± SEM derived from n ≥ 145 cells examined over two independent experiments; *p* values are indicated, unpaired two-tailed t test.

**Extended Data Fig. 4:**
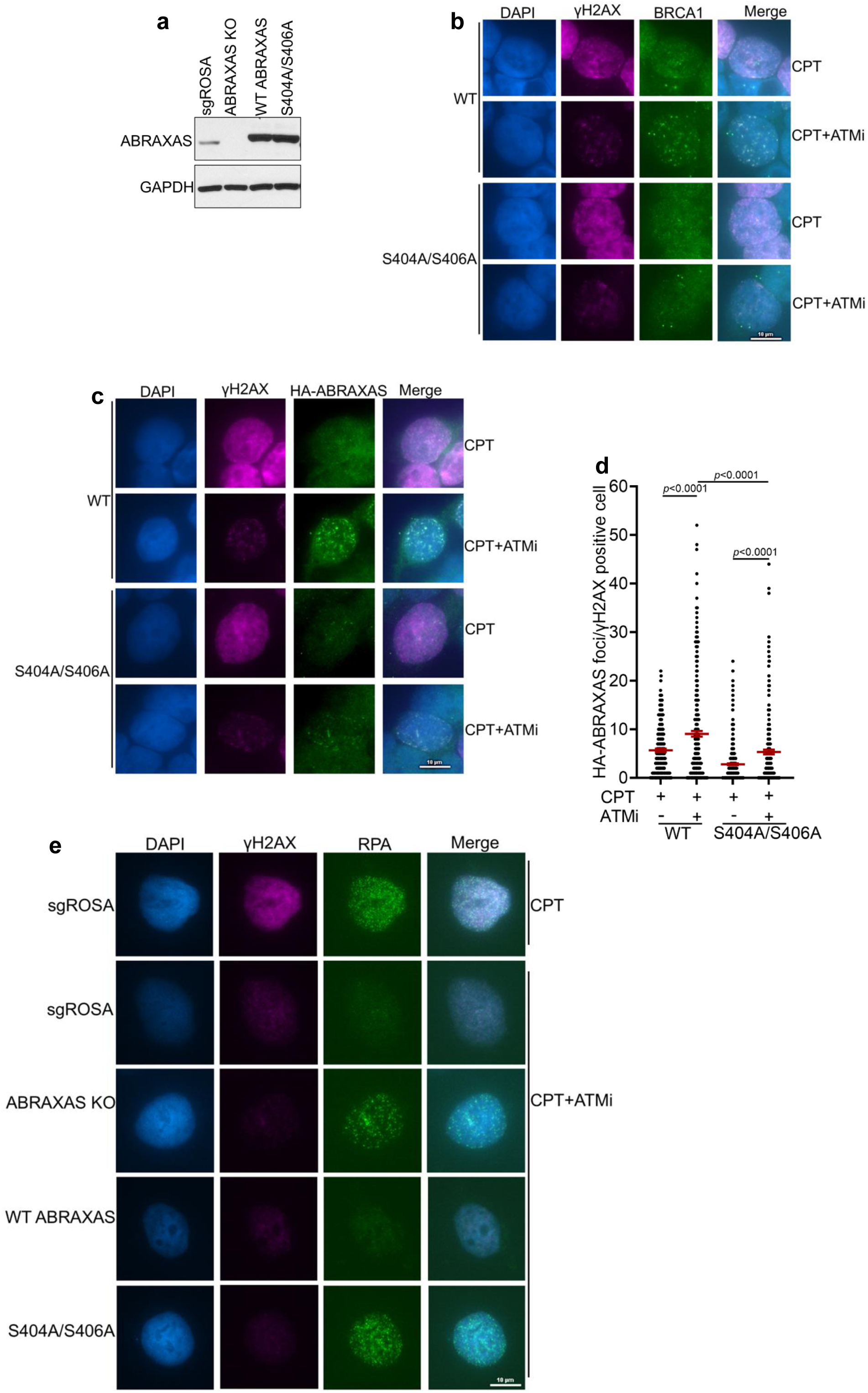
BRCA1 interaction with the A-complex suppresses end resection in ATM inhibited cells. **a**, Western blots showing ABRAXAS levels in ABRAXAS KO HT-29 cells complemented with WT or S404A/S406A ABRAXAS. GAPDH serves as loading control. **b,** Representative immunofluorescence images of BRCA1 foci in WT and S404A/S406A ABRAXAS expressing HT-29 cells exposed to CPT (1 µM) alone or in combination with ATMi (250 nM) for 1 h. ATMi was added 10 min before CPT treatment. γH2AX marks cells with CPT-induced replication damage. **c,** Representative immunofluorescence images of HA-ABRAXAS foci in WT and S404A/S406A ABRAXAS expressing HT-29 cells treated with CPT (1 µM) +/- ATMi (250 nM) for 1 h. **d,** Scatter plot showing quantification of the data as described in (c). Experiments were performed three times with similar results. Data represents mean ± SEM derived from n ≥ 200 cells examined over two independent experiments; *p* values are indicated, unpaired two-tailed t test. **e,** ABRAXAS S404A/S406A mutations restore end resection in ATM-inhibited cells. Representative immunofluorescence images of RPA foci in control (sgROSA), ABRAXAS KO, WT and S404A/S406A ABRAXAS expressing HT-29 cells treated as described in (c). Images were acquired with 100X magnification. Scale bar is shown.

**Extended Data Fig. 5:**
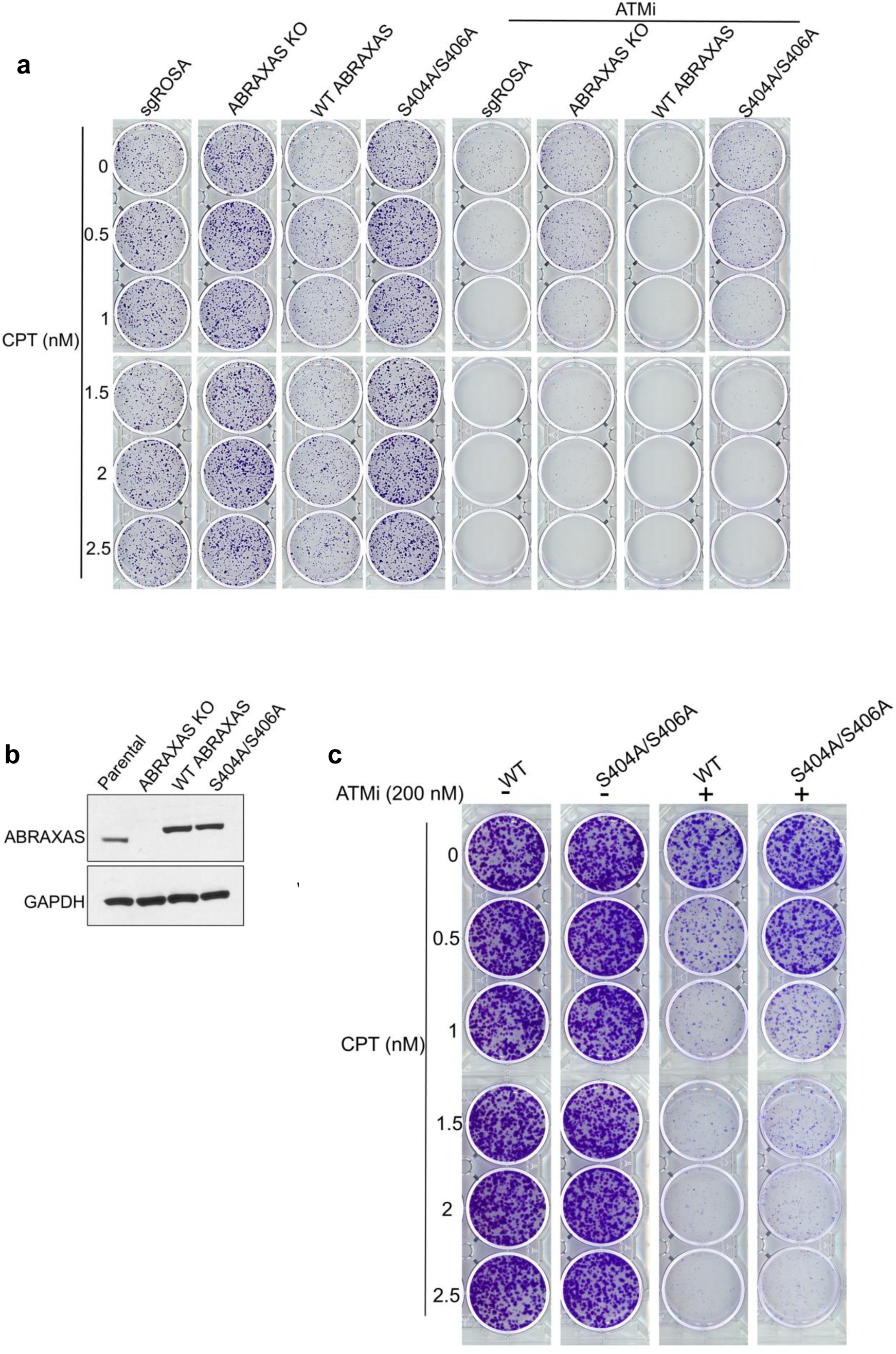
BRCA1 interaction with the A-complex suppresses cell viability upon combined CPT and ATMi treatment. **a,** Clonogenic survival assays showing CPT+ATMi resistance in HT-29 cells expressing ABRAXAS S404A/S406A mutant. **b,** Western blots showing ABRAXAS levels in ABRAXAS KO HEK293T cells complemented with WT or S404A/S406A ABRAXAS. GAPDH serves as loading control. **c,** Clonogenic survival assay showing CPT+ATMi resistance in ABRAXAS S404A/S406A expressing HEK293T cells.

**Extended Data Fig. 6:**
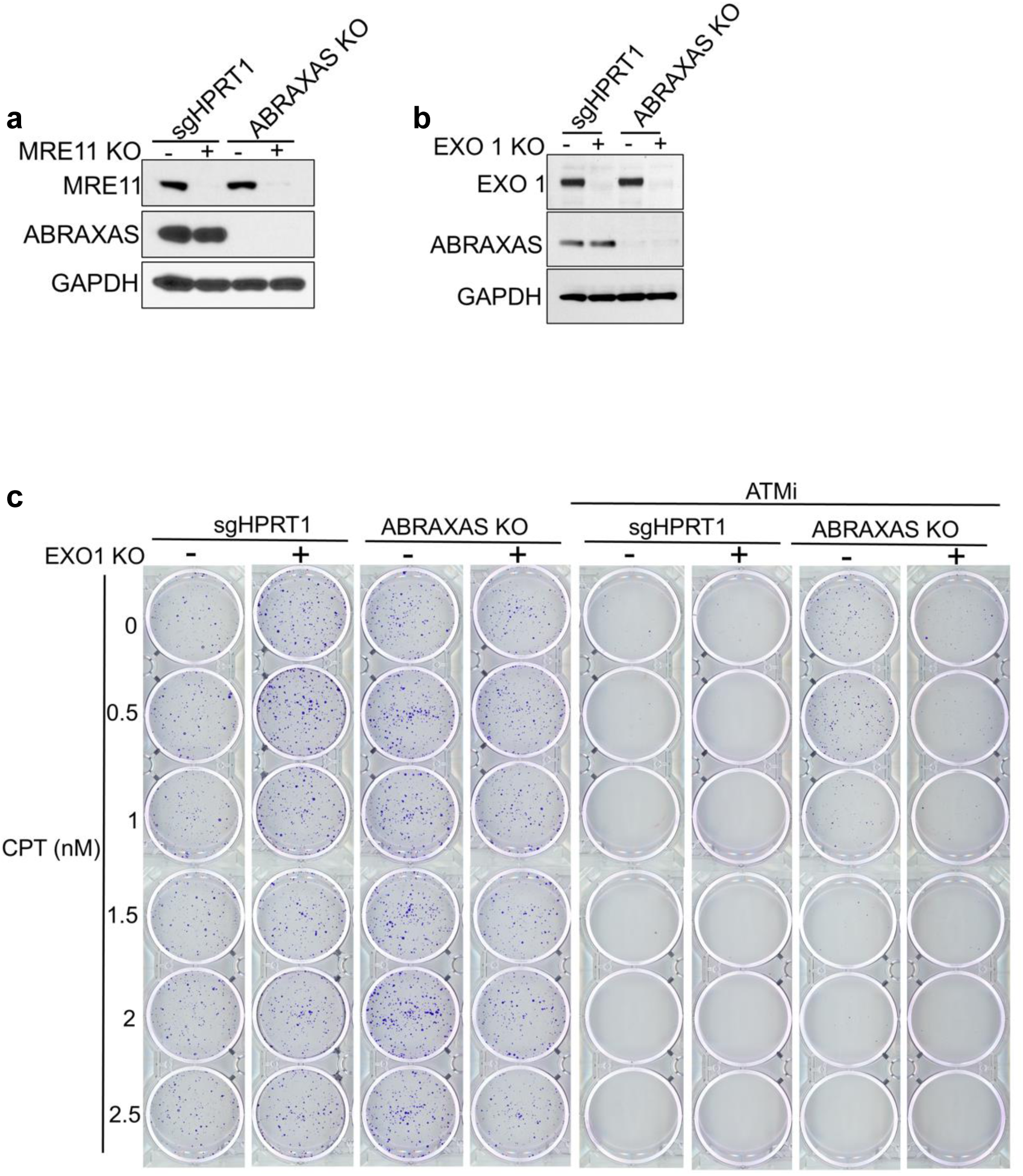
EXO1 KO re-sensitizes ABRAXAS KO cells to combined CPT and ATMi treatment. **(a, b),** Western blots showing MRE11 (a) or EXO1(b) KO in control (sgHPRT1) and ABRAXAS KO HT-29 cells. GAPDH serves as loading control. **c,** Clonogenic survival assays showing increased CPT+ATMi sensitivity in ABRAXAS KO HT-29 cells upon EXO1 KO. Experiments were repeated *n*=3 times with similar results.

**Extended Data Fig. 7:**
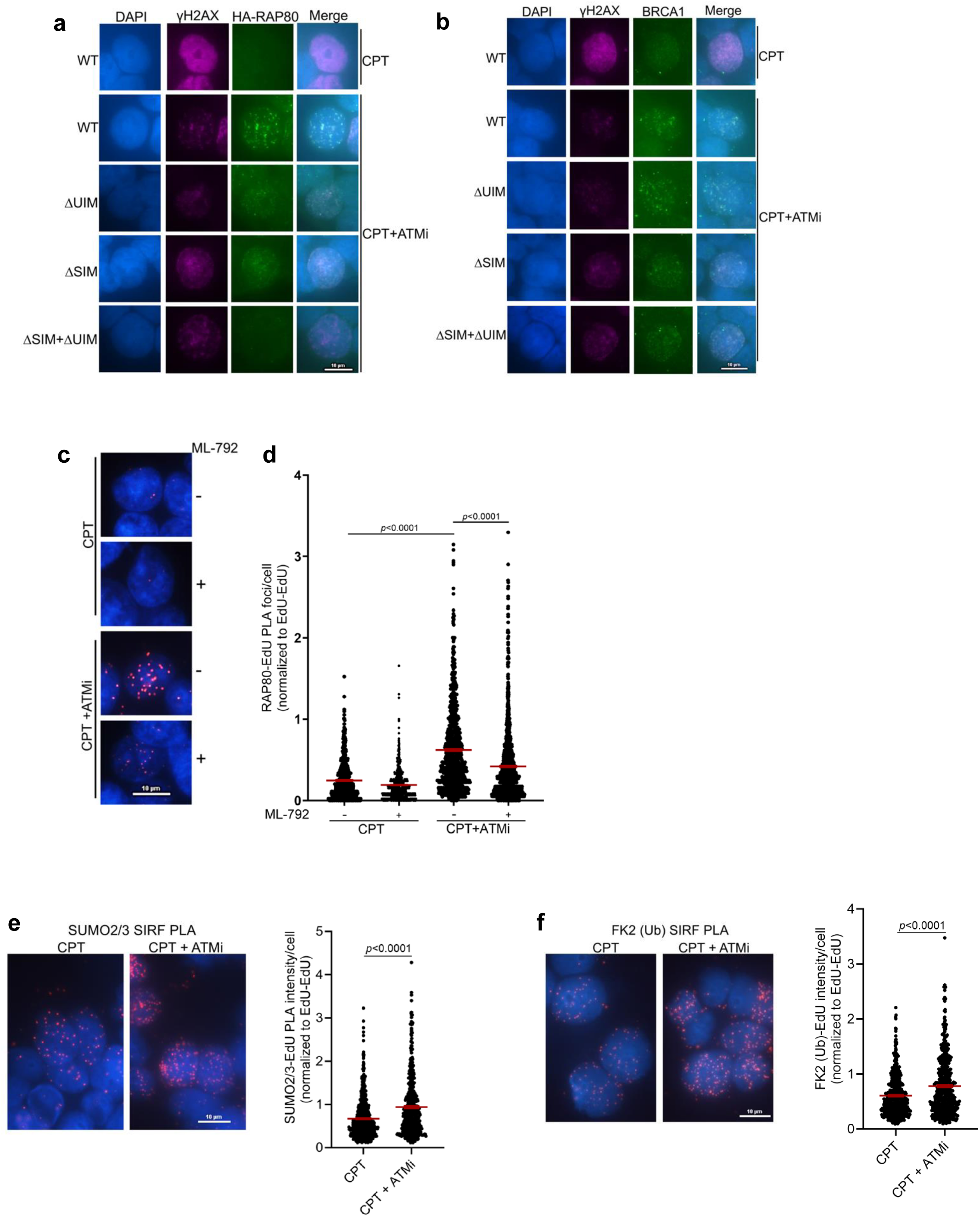
Combined SUMO and ubiquitin recognition by RAP80 facilitates BRCA1-A complex recruitment at damaged chromatin upon ATM inhibition. **a,** Representative immunofluorescence images of HA-RAP80 foci in WT, ΔUIM, ΔSIM or ΔSIM + ΔUIM expressing HT-29 cells treated with CPT (1 µM) +/- ATMi (250 nM) for 1 h. ATMi was added 10 min before CPT treatment. **b,** Representative BRCA1 foci in WT, ΔUIM, ΔSIM or ΔSIM + ΔUIM expressing HT-29 cells treated as described in (a). **(c, d)** Inhibition of SUMOylation by the SUMO E1 inhibitor ML-792 reduces BRCA1-A damage localization in ATMi-treated cells. Representative images of RAP80-EdU PLA in HT-29 cells treated with 50 nM CPT+/- 250 nM ATMi. Cells were optionally treated with ML-792 (1 µM) for 16 h prior to CPT treatment (c). Quantification of RAP80-SIRF PLA foci is shown (d). Data represent mean ± SEM derived from n ≥ 545 cells examined over two independent experiments; *p* values are indicated, unpaired two-tailed t test. **e,** Left panel: PLA-SIRF experiment showing increased SUMO2/3 abundance at CPT-damaged fork upon ATM inhibition. HT-29 cells were optionally pretreated with ATMi (250 nM) for 10 minutes followed by CPT (1 µM) treatment for 40 minutes. EdU was added during the last 20 min of CPT treatment. ATMi was present throughout the experiment. Right panel: Scatter plot showing quantification of SUMO2/3-EdU SIRF foci normalized to average EdU-EdU signal in each experimental condition. Experiments were performed three times with similar results. Data represent mean ± SEM derived from n ≥ 500 cells examined over two independent experiments; *p* values are indicated, unpaired two-tailed t test. **f,** Left panel: PLA-SIRF experiment showing FK2-Ub accumulation at CPT-damaged fork upon ATM inhibition. HT-29 cells were treated as described in (e). Right panel: Quantification of the data is shown. Experiments were performed four times with similar results. Data represent mean ± SEM derived from n ≥ 506 cells examined over two independent experiments; *p* values are indicated, unpaired two-tailed t test.

**Extended Data Fig. 8:**
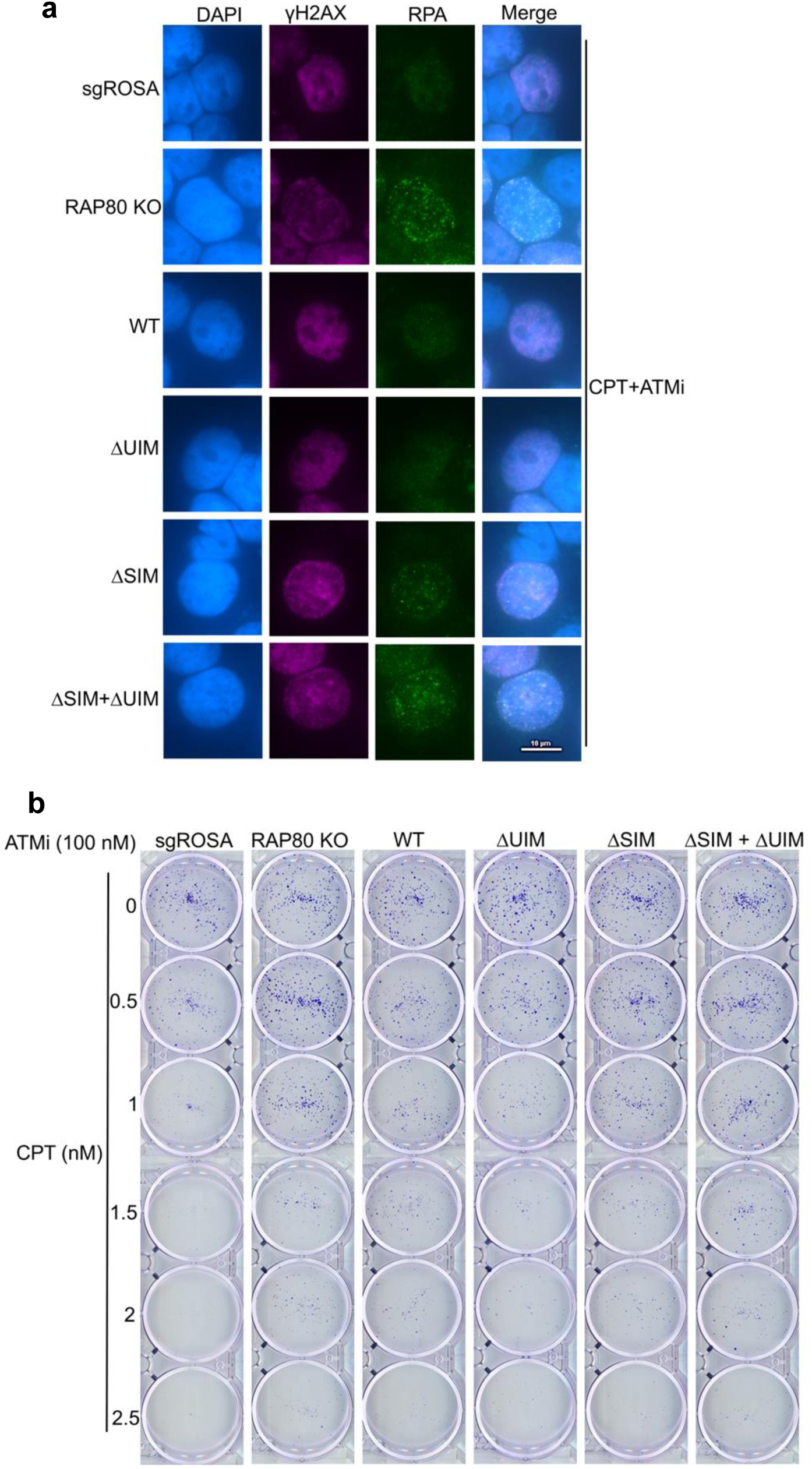
Combined SUMO and ubiquitin recognition by RAP80 suppresses end resection and viability in ATM-inhibited cells. **a,** Representative RPA immunofluorescence in control (sgROSA) and RAP80 KO HT-29 cells complemented with WT or RAP80 deletion mutants. Cells were pretreated with ATMi (250 nM) for 10 min followed by CPT (1 µM) treatment for 1 h. **b,** Representative colony images of control (sgROSA) or RAP80 KO HT-29 cells harboring empty vector (EV) or complemented with WT or RAP80 deletion mutants, Cells were treated with 100 nM ATMi in combination with the indicated doses of CPT.

**Extended Data Fig. 9:**
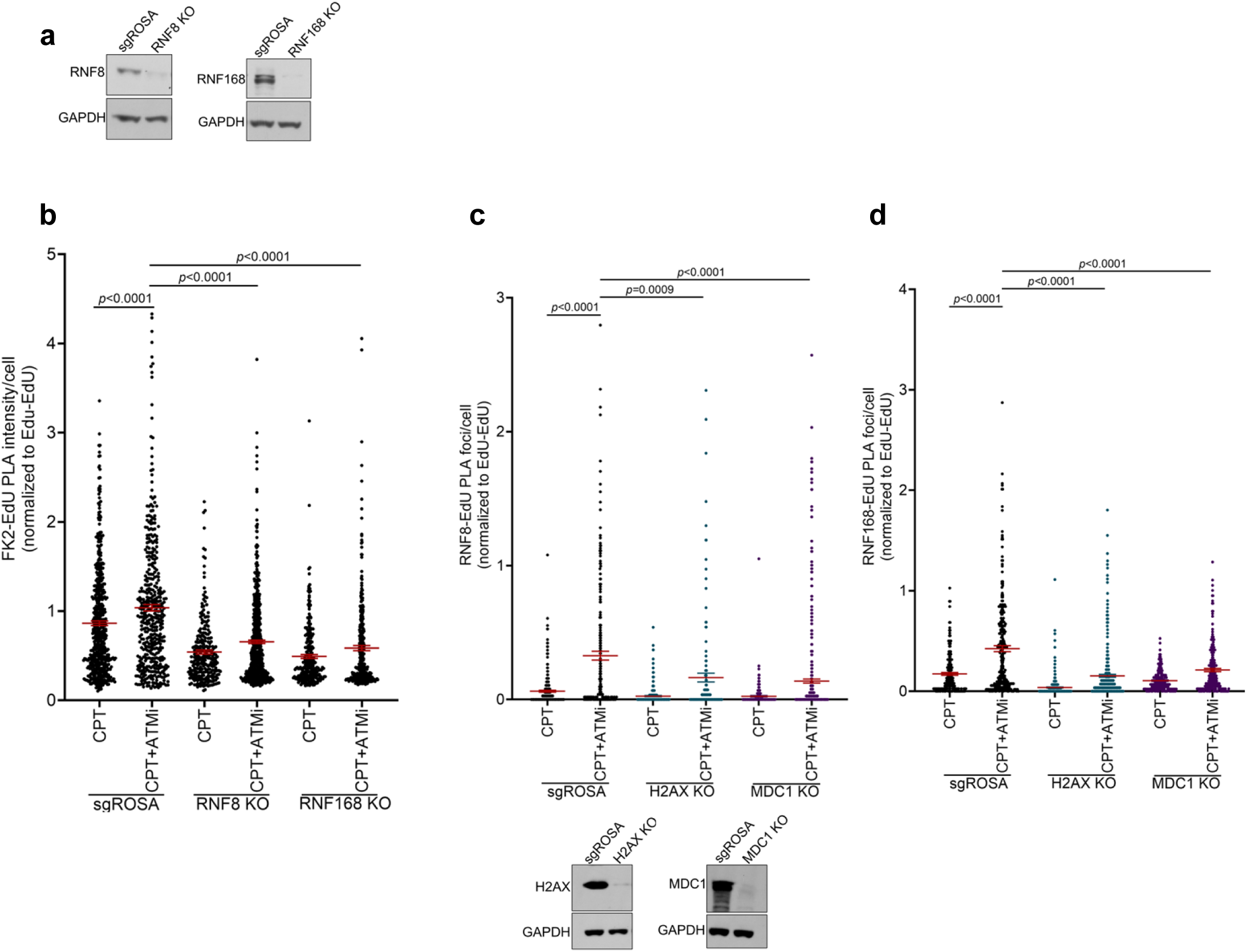
ATM inhibition increases RNF8 and RNF168 dependent ubiquitylation of damaged replication forks. **a,** Western blots showing RNF8 or RNF168 KO in HT-29 cells. GAPDH serves as loading control. **b,** Scatter plot showing quantification of FK2-EdU PLA signal in RNF8 or RNF168 KO HT-29 cells treated as indicated. Data represent mean ± SEM derived from n ≥ 261 cells examined over two independent experiments; *p* values are indicated, unpaired two-tailed t test. **(c, d)** Scatter plot showing quantification of RNF8-EdU (c) and RNF168-EdU (d) PLA signal in H2AX or MDC1 KO HT-29 cells treated as indicated. Data represent mean ± SEM derived from n ≥ 144 (c) and n ≥ 182 (d) cells examined over two independent experiments; *p* values are indicated, unpaired two-tailed t test. H2AX and MDC1 KO in HT-29 cells were verified by western blotting. GAPDH serves as loading control.

**Extended Data Fig. 10:**
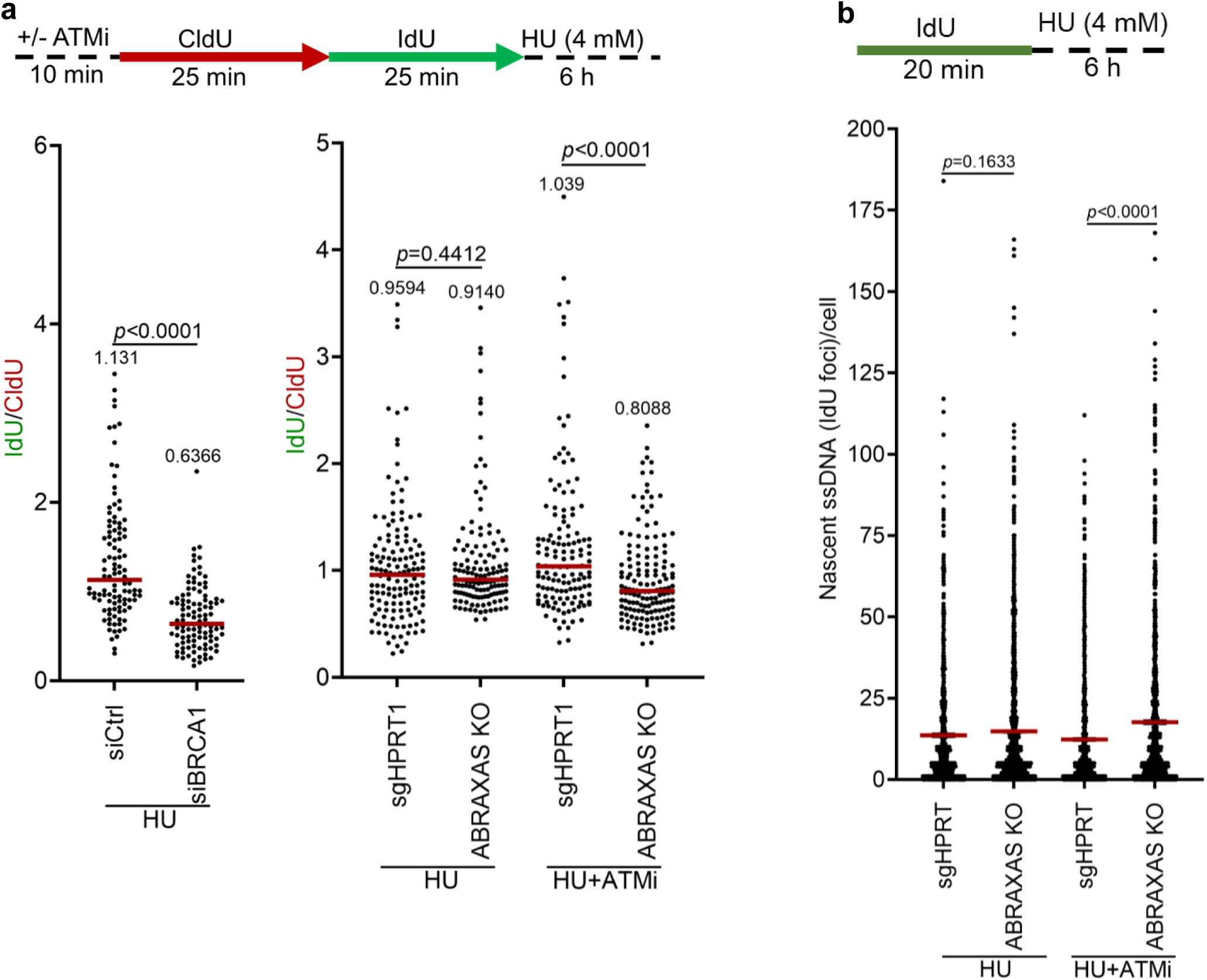
BRCA1-A complex loss causes controlled resection of HU-stalled forks in ATM-inhibited cells. **a,** Fork degradation experiment in BRCA1-depleted or ABRAXAS KO HT-29 cells treated with HU +/- ATMi as indicated. Horizontal red bars indicate median of IdU/CldU ratios. Median IdU/CldU values are indicated. *p* values were derived from n ≥ 100 DNA fibers using two-tailed Mann–Whitney test. **b,** Scatter dot plot showing quantification of native IdU experiment results in control (sgHPRT1) and ABRAXAS KO HT-29 cells treated with HU+/-ATMi as indicated. Data represent mean ± SEM derived from n ≥ 829 cells examined over two independent experiments; *p* values are indicated, unpaired two-tailed t test.

**Extended Data Fig. 11:**
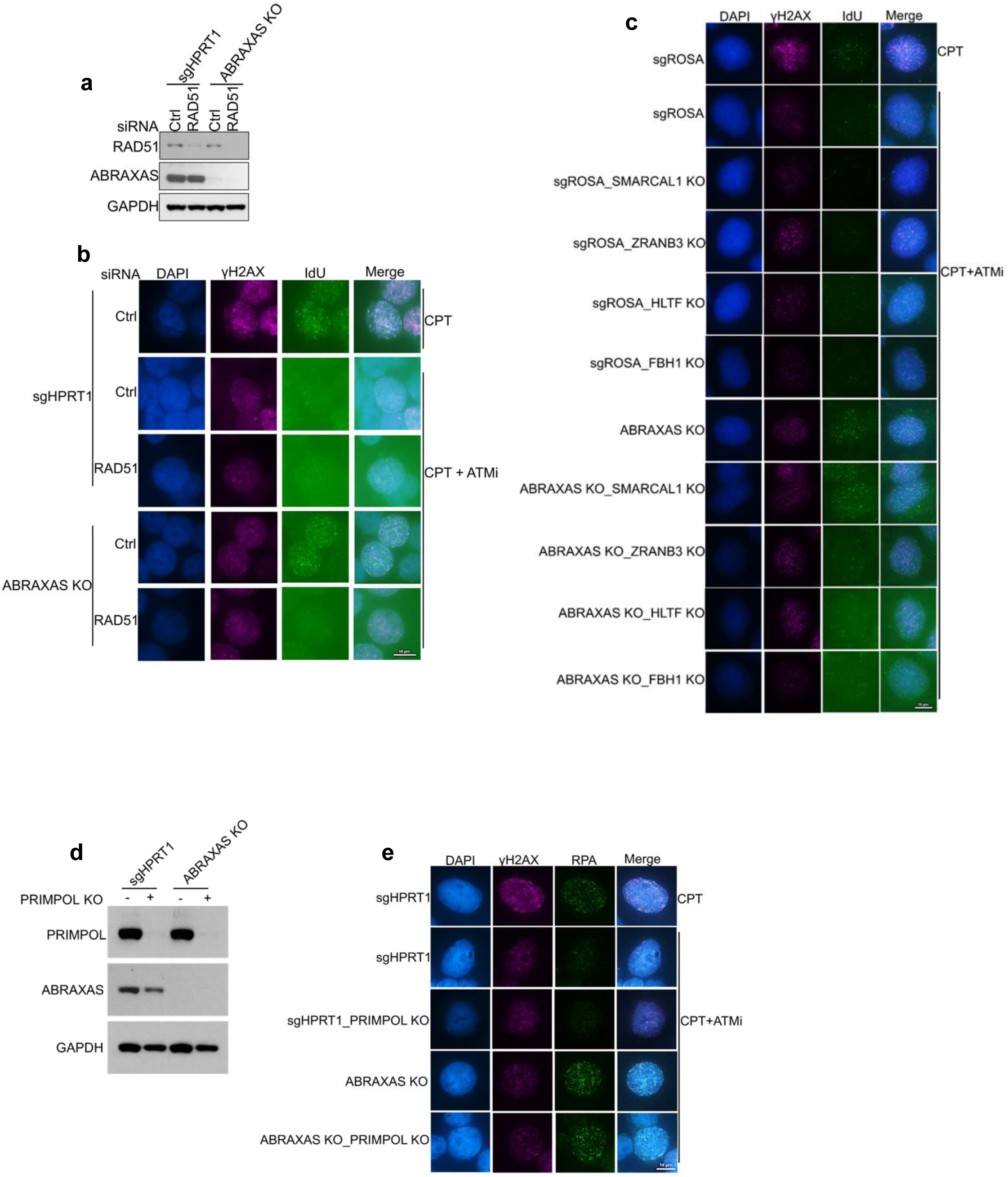
Reversed forks but not ssDNA gaps are resection substrates in BRCA1-A deficient cells. **a,** Western blots showing RAD51 levels in control (sgHPRT1) and ABRAXAS KO HT-29 cells transfected with control or RAD51 siRNA. GAPDH serves as loading control. **b**, Representative images of anti-IdU immunofluorescence detected under non-denaturing conditions in control and ABRAXAS KO cells upon siRNA-mediated depletion of RAD51. Cells were treated with 50 nM CPT +/- 250 nM ATMi for 1 h. **c,** Representative images of anti-IdU immunofluorescence representing ssDNA at reversed forks in U2OS control (sgROSA) and ABRAXAS KO cells upon SMARCAL1, ZRANB3, HLTF or FBH1 KO. Cells were treated with 50 nM CPT for 60 minutes. IdU was added during the last 40 minutes of CPT treatment. Cells were optionally pre-treated with ATMi (250 nM) 10 minutes before CPT addition and continued throughout the experiment. Scale bar is shown. **d,** Western blots showing PRIMPOL KO in control (sgHPRT1) and ABRAXAS KO HT-29 cells. GAPDH serves as a loading control. **e,** Representative RPA immunofluorescence in control (sgHPRT1) and ABRAXAS KO HT-29 cells upon PRIMPOL KO. Cells were treated with 50 nM CPT +/- ATMi (250 nM) as indicated for 1 h. Scale bar is shown.

**Extended Data Fig. 12:**
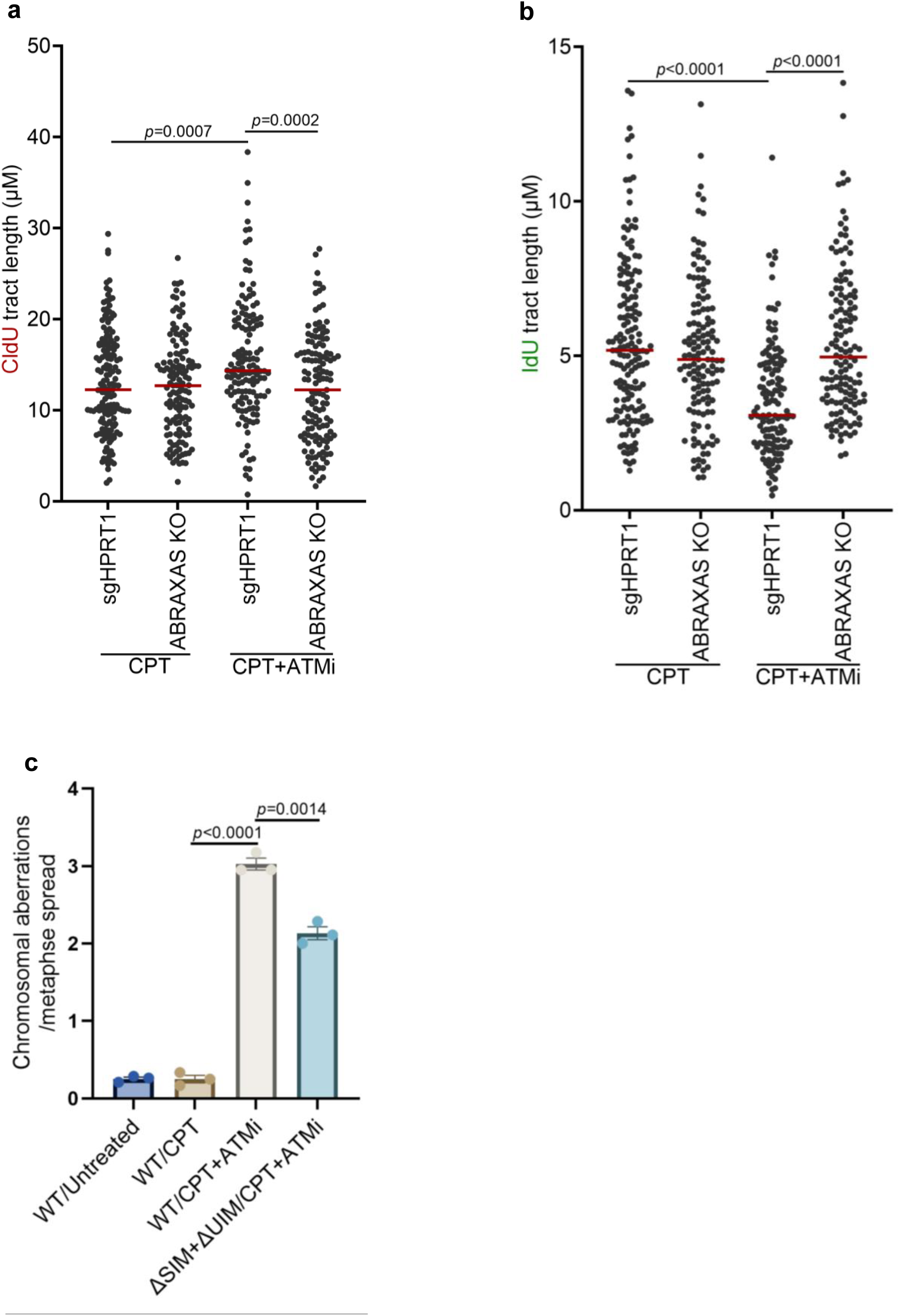
BRCA1-A complex causes fork restart defects and chromosomal aberrations in ATMi-treated cells. **(a,b)** Scatter plot showing CldU (a) and IdU (b) tract length determined from the fork restart experiments as described in Fig.7a. *p*-values were determined from n ≥ 135 DNA fibers using two-tailed Mann–Whitney test. Red bar indicates median tract length. **c,** Bar graph showing quantification of chromosomal aberrations in HT-29 cells expressing either WT or SIM+UIM deleted RAP80. Data represent mean ± SEM of n=3 repeats. At least 60 metaphase spreads were analyzed for each condition across three repeats; *p* values are indicated; unpaired two-tailed t test.

## Notes

### Competing Interest Statement

The authors have declared no competing interest.

## References

1. Chen, C.C., Feng, W., Lim, P.X., Kass, E.M. & Jasin, M. Homology-Directed Repair and the Role of BRCA1, BRCA2, and Related Proteins in Genome Integrity and Cancer. Annu Rev Cancer Biol 2, 313–336 (2018).

2. Chen, C.C. et al. ATM loss leads to synthetic lethality in BRCA1 BRCT mutant mice associated with exacerbated defects in homology-directed repair. Proc Natl Acad Sci U S A 114, 7665–7670 (2017).

3. Scully, R. et al. Genetic analysis of BRCA1 function in a defined tumor cell line. Mol Cell 4, 1093–1099 (1999).

4. Renwick, A. et al. ATM mutations that cause ataxia-telangiectasia are breast cancer susceptibility alleles. Nat Genet 38, 873–875 (2006).

5. Savitsky, K. et al. A single ataxia telangiectasia gene with a product similar to PI-3 kinase. Science 268, 1749–1753 (1995).

6. Wilcox, N. et al. Exome sequencing identifies breast cancer susceptibility genes and defines the contribution of coding variants to breast cancer risk. Nat Genet 55, 1435–1439 (2023).

7. Swift, M., Reitnauer, P.J., Morrell, D. & Chase, C.L. Breast and other cancers in families with ataxia-telangiectasia. N Engl J Med 316, 1289–1294 (1987).

8. Walsh, T. & King, M.C. Ten genes for inherited breast cancer. Cancer Cell 11, 103–105 (2007).

9. Moser, S.C. & Jonkers, J. Thirty Years of BRCA1: Mechanistic Insights and Their Impact on Mutation Carriers. Cancer Discov 15, 461–480 (2025).

10. Nik-Zainal, S. et al. Mutational processes molding the genomes of 21 breast cancers. Cell 149, 979–993 (2012).

11. Ashworth, A. & Lord, C.J. Synthetic lethal therapies for cancer: what’s next after PARP inhibitors? Nat Rev Clin Oncol 15, 564–576 (2018).

12. Kilgas, S., Swift, M.L. & Chowdhury, D. 53BP1-the ‘Pandora’s box’ of genome integrity. DNA Repair (Amst) 144, 103779 (2024).

13. Matsuoka, S. et al. ATM and ATR substrate analysis reveals extensive protein networks responsive to DNA damage. Science 316, 1160–1166 (2007).

14. Sawyer, S.L. et al. Biallelic mutations in BRCA1 cause a new Fanconi anemia subtype. Cancer Discov 5, 135–142 (2015).

15. Jin, Y. et al. Cell cycle-dependent colocalization of BARD1 and BRCA1 proteins in discrete nuclear domains. Proc Natl Acad Sci U S A 94, 12075–12080 (1997).

16. Jiang, Q. & Greenberg, R.A. Deciphering the BRCA1 Tumor Suppressor Network. J Biol Chem 290, 17724–17732 (2015).

17. Greenberg, R.A. et al. Multifactorial contributions to an acute DNA damage response by BRCA1/BARD1-containing complexes. Genes Dev 20, 34–46 (2006).

18. Wang, B. et al. Abraxas and RAP80 form a BRCA1 protein complex required for the DNA damage response. Science 316, 1194–1198 (2007).

19. Kim, H., Chen, J. & Yu, X. Ubiquitin-binding protein RAP80 mediates BRCA1-dependent DNA damage response. Science 316, 1202–1205 (2007).

20. Sobhian, B. et al. RAP80 targets BRCA1 to specific ubiquitin structures at DNA damage sites. Science 316, 1198–1202 (2007).

21. Sy, S.M., Huen, M.S. & Chen, J. PALB2 is an integral component of the BRCA complex required for homologous recombination repair. Proc Natl Acad Sci U S A 106, 7155–7160 (2009).

22. Zhang, F. et al. PALB2 links BRCA1 and BRCA2 in the DNA-damage response. Curr Biol 19, 524–529 (2009).

23. Yu, X., Chini, C.C., He, M., Mer, G. & Chen, J. The BRCT domain is a phospho-protein binding domain. Science 302, 639–642 (2003).

24. Manke, I.A., Lowery, D.M., Nguyen, A. & Yaffe, M.B. BRCT repeats as phosphopeptide-binding modules involved in protein targeting. Science 302, 636–639 (2003).

25. Reczek, C.R., Szabolcs, M., Stark, J.M., Ludwig, T. & Baer, R. The interaction between CtIP and BRCA1 is not essential for resection-mediated DNA repair or tumor suppression. J Cell Biol 201, 693–707 (2013).

26. Polato, F. et al. CtIP-mediated resection is essential for viability and can operate independently of BRCA1. J Exp Med 211, 1027–1036 (2014).

27. Bridge, W.L., Vandenberg, C.J., Franklin, R.J. & Hiom, K. The BRIP1 helicase functions independently of BRCA1 in the Fanconi anemia pathway for DNA crosslink repair. Nat Genet 37, 953–957 (2005).

28. Guzzo, C.M. et al. RNF4-dependent hybrid SUMO-ubiquitin chains are signals for RAP80 and thereby mediate the recruitment of BRCA1 to sites of DNA damage. Sci Signal 5, ra88 (2012).

29. Nakamura, K. et al. H4K20me0 recognition by BRCA1-BARD1 directs homologous recombination to sister chromatids. Nat Cell Biol 21, 311–318 (2019).

30. Becker, J.R. et al. BARD1 reads H2A lysine 15 ubiquitination to direct homologous recombination. Nature 596, 433–437 (2021).

31. Burdett, H. et al. BRCA1-BARD1 combines multiple chromatin recognition modules to bridge nascent nucleosomes. Nucleic Acids Res 51, 11080–11103 (2023).

32. Witus, S.R. et al. BRCA1/BARD1 intrinsically disordered regions facilitate chromatin recruitment and ubiquitylation. EMBO J 42, e113565 (2023).

33. Hu, Q. et al. Mechanisms of BRCA1-BARD1 nucleosome recognition and ubiquitylation. Nature 596, 438–443 (2021).

34. Shakya, R. et al. BRCA1 tumor suppression depends on BRCT phosphoprotein binding, but not its E3 ligase activity. Science 334, 525–528 (2011).

35. Jiang, Q. et al. MERIT40 cooperates with BRCA2 to resolve DNA interstrand cross-links. Genes Dev 29, 1955–1968 (2015).

36. Wu, J., Liu, C., Chen, J. & Yu, X. RAP80 protein is important for genomic stability and is required for stabilizing BRCA1-A complex at DNA damage sites in vivo. J Biol Chem 287, 22919–22926 (2012).

37. Coleman, K.A. & Greenberg, R.A. The BRCA1-RAP80 complex regulates DNA repair mechanism utilization by restricting end resection. J Biol Chem 286, 13669–13680 (2011).

38. Hu, Y. et al. RAP80-directed tuning of BRCA1 homologous recombination function at ionizing radiation-induced nuclear foci. Genes Dev 25, 685–700 (2011).

39. Balmus, G. et al. ATM orchestrates the DNA-damage response to counter toxic non-homologous end-joining at broken replication forks. Nat Commun 10, 87 (2019).

40. Cai, M.Y. et al. Cooperation of the ATM and Fanconi Anemia/BRCA Pathways in Double-Strand Break End Resection. Cell Rep 30, 2402–2415.e2405 (2020).

41. Holohan, C., Van Schaeybroeck, S., Longley, D.B. & Johnston, P.G. Cancer drug resistance: an evolving paradigm. Nat Rev Cancer 13, 714–726 (2013).

42. Riches, L.C. et al. Pharmacology of the ATM Inhibitor AZD0156: Potentiation of Irradiation and Olaparib Responses Preclinically. Mol Cancer Ther 19, 13–25 (2020).

43. Nakamura, K. et al. Proteome dynamics at broken replication forks reveal a distinct ATM-directed repair response suppressing DNA double-strand break ubiquitination. Mol Cell 81, 1084–1099.e1086 (2021).

44. Bunting, S.F. et al. 53BP1 inhibits homologous recombination in Brca1-deficient cells by blocking resection of DNA breaks. Cell 141, 243–254 (2010).

45. Bouwman, P. et al. 53BP1 loss rescues BRCA1 deficiency and is associated with triple-negative and BRCA-mutated breast cancers. Nat Struct Mol Biol 17, 688–695 (2010).

46. Ray Chaudhuri, A., et al. Topoisomerase I poisoning results in PARP-mediated replication fork reversal. Nat Struct Mol Biol 19, 417–423 (2012).

47. Zellweger, R. et al. Rad51-mediated replication fork reversal is a global response to genotoxic treatments in human cells. J Cell Biol 208, 563–579 (2015).

48. Lemaçon, D. et al. MRE11 and EXO1 nucleases degrade reversed forks and elicit MUS81-dependent fork rescue in BRCA2-deficient cells. Nature Communications 8, 860 (2017).

49. Thangavel, S. et al. DNA2 drives processing and restart of reversed replication forks in human cells. J Cell Biol 208, 545–562 (2015).

50. Bhat, K.P. & Cortez, D. RPA and RAD51: fork reversal, fork protection, and genome stability. Nat Struct Mol Biol 25, 446–453 (2018).

51. Schlacher, K. et al. Double-Strand Break Repair-Independent Role for BRCA2 in Blocking Stalled Replication Fork Degradation by MRE11. Cell 145, 993 (2011).

52. Couch, F.B. et al. ATR phosphorylates SMARCAL1 to prevent replication fork collapse. Genes Dev 27, 1610–1623 (2013).

53. Fugger, K. et al. FBH1 Catalyzes Regression of Stalled Replication Forks. Cell Rep 10, 1749–1757 (2015).

54. Iannascoli, C., Palermo, V., Murfuni, I., Franchitto, A. & Pichierri, P. The WRN exonuclease domain protects nascent strands from pathological MRE11/EXO1-dependent degradation. Nucleic Acids Res 43, 9788–9803 (2015).

55. Wu, Q. et al. Structure of BRCA1-BRCT/Abraxas Complex Reveals Phosphorylation-Dependent BRCT Dimerization at DNA Damage Sites. Mol Cell 61, 434–448 (2016).

56. Adkins, N.L., Niu, H., Sung, P. & Peterson, C.L. Nucleosome dynamics regulates DNA processing. Nat Struct Mol Biol 20, 836–842 (2013).

57. Clouaire, T. & Legube, G. A Snapshot on *Cis* Chromatin Response to DNA Double-Strand Breaks. Trends in Genetics 35, 330–345 (2019).

58. Hamiche, A., Sandaltzopoulos, R., Gdula, D.A. & Wu, C. ATP-Dependent Histone Octamer Sliding Mediated by the Chromatin Remodeling Complex NURF. Cell 97, 833–842 (1999).

59. Toiber, D. et al. SIRT6 Recruits SNF2H to DNA Break Sites, Preventing Genomic Instability through Chromatin Remodeling. Molecular Cell 51, 454–468 (2013).

60. Chen, X. et al. The Fun30 nucleosome remodeller promotes resection of DNA double-strand break ends. Nature 489, 576–580 (2012).

61. Delamarre, A. et al. MRX Increases Chromatin Accessibility at Stalled Replication Forks to Promote Nascent DNA Resection and Cohesin Loading. Molecular Cell 77, 395–410.e393 (2020).

62. Ziv, Y. et al. Chromatin relaxation in response to DNA double-strand breaks is modulated by a novel ATM- and KAP-1 dependent pathway. Nat Cell Biol 8, 870–876 (2006).

63. Berkovich, E., Monnat, R.J., Jr. & Kastan, M.B. Roles of ATM and NBS1 in chromatin structure modulation and DNA double-strand break repair. Nat Cell Biol 9, 683–690 (2007).

64. Gaggioli, V. et al. Dynamic de novo heterochromatin assembly and disassembly at replication forks ensures fork stability. Nat Cell Biol 25, 1017–1032 (2023).

65. Trenz, K., Smith, E., Smith, S. & Costanzo, V. ATM and ATR promote Mre11 dependent restart of collapsed replication forks and prevent accumulation of DNA breaks. Embo j 25, 1764–1774 (2006).

66. Jackson, S.P. & Durocher, D. Regulation of DNA damage responses by ubiquitin and SUMO. Mol Cell 49, 795–807 (2013).

67. Liu, W. et al. RAD51 bypasses the CMG helicase to promote replication fork reversal. Science 380, 382–387 (2023).

68. Adolph, M.B. & Cortez, D. Mechanisms and regulation of replication fork reversal. DNA Repair (Amst) 141, 103731 (2024).

69. Vujanovic, M. et al. Replication Fork Slowing and Reversal upon DNA Damage Require PCNA Polyubiquitination and ZRANB3 DNA Translocase Activity. Mol Cell 67, 882–890.e885 (2017).

70. Berti, M. et al. Sequential role of RAD51 paralog complexes in replication fork remodeling and restart. Nature Communications 11, 3531 (2020).

71. Quinet, A. et al. PRIMPOL-Mediated Adaptive Response Suppresses Replication Fork Reversal in BRCA-Deficient Cells. Mol Cell 77, 461–474.e469 (2020).

72. Taglialatela, A. et al. REV1-Polζ maintains the viability of homologous recombination-deficient cancer cells through mutagenic repair of PRIMPOL-dependent ssDNA gaps. Mol Cell 81, 4008–4025.e4007 (2021).

73. Panzarino, N.J. et al. Replication Gaps Underlie BRCA Deficiency and Therapy Response. Cancer Res 81, 1388–1397 (2021).

74. Tirman, S. et al. Temporally distinct post-replicative repair mechanisms fill PRIMPOL-dependent ssDNA gaps in human cells. Mol Cell 81, 4026–4040.e4028 (2021).

75. Hale, A., Dhoonmoon, A., Straka, J., Nicolae, C.M. & Moldovan, G.-L. Multi-step processing of replication stress-derived nascent strand DNA gaps by MRE11 and EXO1 nucleases. Nature Communications 14, 6265 (2023).

76. Petermann, E., Orta, M.L., Issaeva, N., Schultz, N. & Helleday, T. Hydroxyurea-stalled replication forks become progressively inactivated and require two different RAD51-mediated pathways for restart and repair. Mol Cell 37, 492–502 (2010).

77. Verma, P. et al. ALC1 links chromatin accessibility to PARP inhibitor response in homologous recombination-deficient cells. Nat Cell Biol 23, 160–171 (2021).

78. Shiloh, Y. & Ziv, Y. The ATM protein kinase: regulating the cellular response to genotoxic stress, and more. Nat Rev Mol Cell Biol 14, 197–210 (2013).

79. Yamamoto, K. et al. Kinase-dead ATM protein is highly oncogenic and can be preferentially targeted by Topo-isomerase I inhibitors. Elife 5 (2016).

80. Choi, M., Kipps, T. & Kurzrock, R. ATM Mutations in Cancer: Therapeutic Implications. Mol Cancer Ther 15, 1781–1791 (2016).

81. Smith, P.J., Makinson, T.A. & Watson, J.V. Enhanced sensitivity to camptothecin in ataxia-telangiectasia cells and its relationship with the expression of DNA topoisomerase I. Int J Radiat Biol 55, 217–231 (1989).

82. Wang, C. et al. Genetic vulnerabilities upon inhibition of DNA damage response. Nucleic Acids Res 49, 8214–8231 (2021).

83. Cai, M.-Y. et al. Cooperation of the ATM and Fanconi Anemia/BRCA Pathways in Double-Strand Break End Resection. Cell Reports 30, 2402–2415.e2405 (2020).

84. Jiang, Q. et al. Autologous K63 deubiquitylation within the BRCA1-A complex licenses DNA damage recognition. J Cell Biol 221 (2022).

85. Goodarzi, A.A. et al. ATM Signaling Facilitates Repair of DNA Double-Strand Breaks Associated with Heterochromatin. Molecular Cell 31, 167–177 (2008).

86. Sherker, A. et al. Two redundant ubiquitin-dependent pathways of BRCA1 localization to DNA damage sites. EMBO reports 22, e53679 (2021).

87. Zhu, C. et al. Profilin-1 regulates DNA replication forks in a context-dependent fashion by interacting with SNF2H and BOD1L. Nat Commun 13, 6531 (2022).

88. García-Rodríguez, N., Domínguez-García, I., Domínguez-Pérez, M. del C. & Huertas, P. EXO1 and DNA2-mediated ssDNA gap expansion is essential for ATR activation and to maintain viability in BRCA1-deficient cells. Nucleic Acids Research 52, 6376–6391 (2024).

89. van de Kooij, B. et al. EXO1 protects BRCA1-deficient cells against toxic DNA lesions. Mol Cell 84, 659–674.e657 (2024).

90. Sartori, A.A. et al. Human CtIP promotes DNA end resection. Nature 450, 509–514 (2007).

91. Yan, J. et al. The ubiquitin-interacting motif containing protein RAP80 interacts with BRCA1 and functions in DNA damage repair response. Cancer Res 67, 6647–6656 (2007).

92. Cortez, D., Wang, Y., Qin, J. & Elledge, S.J. Requirement of ATM-dependent phosphorylation of brca1 in the DNA damage response to double-strand breaks. Science 286, 1162–1166 (1999).

93. Gatei, M. et al. Ataxia telangiectasia mutated (ATM) kinase and ATM and Rad3 related kinase mediate phosphorylation of Brca1 at distinct and overlapping sites. In vivo assessment using phospho-specific antibodies. J Biol Chem 276, 17276–17280 (2001).

94. Ryu, H.Y. & Hochstrasser, M. Histone sumoylation and chromatin dynamics. Nucleic Acids Res 49, 6043–6052 (2021).

95. García-Rodríguez, N., Wong, R.P. & Ulrich, H.D. Functions of Ubiquitin and SUMO in DNA Replication and Replication Stress. Front Genet 7, 87 (2016).

96. Garvin, A.J. et al. SUMO4 promotes SUMO deconjugation required for DNA double-strand-break repair. Mol Cell 85, 877–893.e879 (2025).

97. Quinet, A., Lemaçon, D. & Vindigni, A. Replication Fork Reversal: Players and Guardians. Mol Cell 68, 830–833 (2017).

98. Neelsen, K.J. & Lopes, M. Replication fork reversal in eukaryotes: from dead end to dynamic response. Nature Reviews Molecular Cell Biology 16, 207–220 (2015).

99. Schlacher, K., Wu, H. & Jasin, M. A distinct replication fork protection pathway connects Fanconi anemia tumor suppressors to RAD51-BRCA1/2. Cancer cell 22, 106–116 (2012).

100. Liu, W., Krishnamoorthy, A., Zhao, R. & Cortez, D. Two replication fork remodeling pathways generate nuclease substrates for distinct fork protection factors. Science Advances 6, eabc3598.

101. Sogo, J.M., Lopes, M. & Foiani, M. Fork Reversal and ssDNA Accumulation at Stalled Replication Forks Owing to Checkpoint Defects. Science 297, 599–602 (2002).

102. Scully, R., Walter, J.C. & Nussenzweig, A. One-ended and two-ended breaks at nickase-broken replication forks. DNA Repair 144, 103783 (2024).

103. Bai, G. et al. HLTF Promotes Fork Reversal, Limiting Replication Stress Resistance and Preventing Multiple Mechanisms of Unrestrained DNA Synthesis. Mol Cell 78, 1237–1251.e1237 (2020).

104. Taglialatela, A. et al. Restoration of Replication Fork Stability in BRCA1- and BRCA2-Deficient Cells by Inactivation of SNF2-Family Fork Remodelers. Mol Cell 68, 414–430.e418 (2017).

105. Fugger, K. et al. FBH1 Catalyzes Regression of Stalled Replication Forks. Cell Reports 10, 1749–1757 (2015).

106. Kolinjivadi, A.M. et al. Smarcal1-Mediated Fork Reversal Triggers Mre11-Dependent Degradation of Nascent DNA in the Absence of Brca2 and Stable Rad51 Nucleofilaments. Mol Cell 67, 867–881.e867 (2017).

107. Bryant, H.E. et al. PARP is activated at stalled forks to mediate Mre11-dependent replication restart and recombination. Embo j 28, 2601–2615 (2009).

108. Ray Chaudhuri, A., et al. Replication fork stability confers chemoresistance in BRCA-deficient cells. Nature 535, 382–387 (2016).

109. Zhang, T. et al. Break-induced replication orchestrates resection-dependent template switching. Nature 619, 201–208 (2023).

110. Ran, F.A. et al. Genome engineering using the CRISPR-Cas9 system. Nat Protoc 8, 2281–2308 (2013).

111. Patterson-Fortin, J., Shao, G., Bretscher, H., Messick, T.E. & Greenberg, R.A. Differential regulation of JAMM domain deubiquitinating enzyme activity within the RAP80 complex. J Biol Chem 285, 30971–30981 (2010).

112. Sirbu, B.M., Couch, F.B. & Cortez, D. Monitoring the spatiotemporal dynamics of proteins at replication forks and in assembled chromatin using isolation of proteins on nascent DNA. Nat Protoc 7, 594–605 (2012).

113. Rappsilber, J., Ishihama, Y. & Mann, M. Stop and go extraction tips for matrix-assisted laser desorption/ionization, nanoelectrospray, and LC/MS sample pretreatment in proteomics. Anal Chem 75, 663–670 (2003).

114. Peng, J. & Gygi, S.P. Proteomics: the move to mixtures. J Mass Spectrom 36, 1083–1091 (2001).

115. Eng, J.K., McCormack, A.L. & Yates, J.R. An approach to correlate tandem mass spectral data of peptides with amino acid sequences in a protein database. J Am Soc Mass Spectrom 5, 976–989 (1994).

116. Garcia-Moreno, A. et al. Functional Enrichment Analysis of Regulatory Elements. Biomedicines 10 (2022).

117. Stirling, D.R. et al. CellProfiler 4: improvements in speed, utility and usability. BMC Bioinformatics 22, 433 (2021).

118. Roy, S. & Schlacher, K. SIRF: A Single-cell Assay for in situ Protein Interaction with Nascent DNA Replication Forks. Bio Protoc 9, e3377 (2019).

119. Roy, S., Luzwick, J.W. & Schlacher, K. SIRF: Quantitative in situ analysis of protein interactions at DNA replication forks. Journal of Cell Biology 217, 1521–1536 (2018).

120. Lopes, M. Electron Microscopy Methods for Studying In Vivo DNA Replication Intermediates, in DNA Replication: Methods and Protocols. (eds. S. Vengrova & J.Z. Dalgaard) 605–631 (Humana Press, Totowa, NJ; 2009).

121. Neelsen, K.J., Chaudhuri, A.R., Follonier, C., Herrador, R. & Lopes, M. Visualization and interpretation of eukaryotic DNA replication intermediates in vivo by electron microscopy. Methods Mol Biol 1094, 177–208 (2014).

122. Schneider, C.A., Rasband, W.S. & Eliceiri, K.W. NIH Image to ImageJ: 25 years of image analysis. Nature Methods 9, 671–675 (2012).

